# Modelling Anaerobic Co-Digestion with Agricultural Feedstock: Model Validation and Cross-Reactor Verification

**DOI:** 10.64898/2026.04.27.721061

**Authors:** Rohit Murali, Benaissa Dekhici, Tao Chen, Dongda Zhang, Michael Short

**Affiliations:** School of Chemistry and Chemical Engineering, University of Surrey, Guildford, GU2 7XH, UK; Surrey Institute for People-Centred Artificial Intelligence, University of Surrey, Guildford, GU2 7XH, UK; School of Chemical Engineering and Analytical Science, The University of Manchester, Manchester, M13 9PL, UK; Supergen Bioenergy Impact Hub, Energy and Bioproducts Research Institute, UK

**Keywords:** Anaerobic Digestion, ADM1, Physical Twin, Co-digestion

## Abstract

As the United Kingdom (UK) targets net-zero emissions by 2050, anaerobic digestion (AD) has become a cornerstone of renewable energy infrastructure. However, mathematical models, such as the Anaerobic Digestion Model No. 1 (ADM1), often struggle with high-solids agricultural feedstocks because they rely on Chemical Oxygen Demand (COD), a metric that introduces significant experimental error. To overcome this, this study applies an established mass-based ADM1 framework tailored for the co-digestion of maize silage and cow manure sourced from a UK AD site. This study uses a parallel reactor framework, using two identical laboratory-scale reactors to physically replicate the dynamic conditions of the full-scale site. A Global Sensitivity Analysis was first conducted, identifying biomass decay and carbohydrate breakdown rates as the most influential factors affecting system stability and model accuracy. The model was calibrated using data from the first reactor and then tested against an independent second reactor subjected to significant organic loading stress. Results show high predictive capabilities, with the model achieving a R^2^ of 0.81 for biogas production during calibration. The model maintained high predictive accuracy during the validation test of the second physical twin, achieving an R^2^ of 0.85, proving that the framework is robust and not overfitted to a single dataset. While predicting rapid fluctuations in pH and alkalinity remains challenging, the mass-based approach effectively forecasts gas yields and process stability. This methodology provides a reliable foundation for robust process modelling, offering a scalable tool for the UK biogas sector to optimise AD.

**Graphical Abstract:** 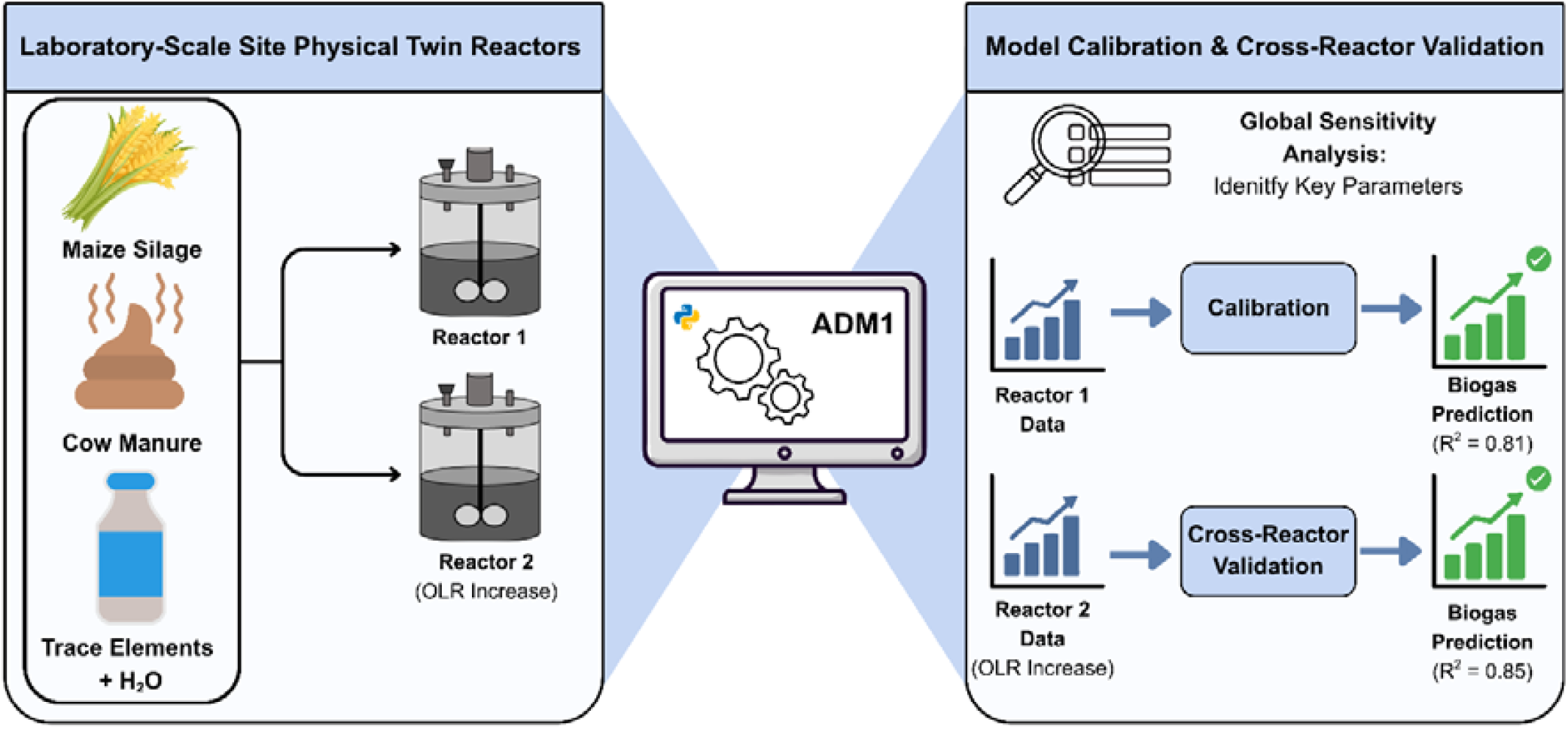

## 1.0 Introduction

### 1.1 The Strategic Imperative of Anaerobic Digestion in the UK

The United Kingdom (UK) is at a pivotal moment in its shift to a low-carbon circular economy. As the nation attempts to achieve its net-zero targets by 2050, the decarbonisation of the energy sector has emerged as a critical policy priority [1]. Within this landscape, AD has evolved from its traditional role as a waste management strategy to become an important aspect of renewable energy infrastructure [2]. The ability of AD to convert organic matter into biogas (a mixture primarily of methane and carbon dioxide) offers a dual benefit as it mitigates fugitive methane emissions from agricultural residues and produces a renewable gas that can be upgraded to biomethane for injection into the national gas grid [2]. To support this, the UK government launched the Green Gas Support Scheme (GGSS) in 2021. The scheme incentivises biomethane injection through tariffs that favour sustainable feedstocks, such as agricultural feedstock, while imposing strict sustainability criteria on purpose-grown energy crops such as maize [2]. Consequently, the operational standard of UK AD plants has shifted towards anaerobic codigestion. This approach, which blends two or more feedstocks, helps improve gas output and process performance while minimising process inhibition and digester stability [3].

However, the biological complexity of AD makes the process naturally sensitive to process disturbances. The sequential degradation of organic substrates by a consortium of hydrolytic, acidogenic, acetogenic, and methanogenic microorganisms requires a delicate balance of intermediate metabolites [4]. In the context of the GGSS, where operators are incentivised to maximise gas product, the drive for higher output creates a significant risk of system failure caused by volatile fatty acid (VFA) accumulation and reactor acidification [5]. This occurs because high organic loading causes fast-growing acidogenic bacteria to produce VFAs at a rate that exceeds the consumption capacity of the slower-growing methanogenic archaea, reducing the system’s buffering capacity [6]. This operational dilemma requires reliable predictive mathematical models that can simulate the dynamic behaviour of the digester under varying organic loading rates (OLR) and feedstock compositions [7].

### 1.2 Background to Process Modelling of AD

Mathematical modelling can provide a framework for understanding and predicting the complicated network of biochemical interactions within a digester. The Anaerobic Digestion Model No. 1 (ADM1), developed by the International Water Association (IWA) Task Group, is the standard model for simulating AD processes [6,8]. The application of ADM1 to the specific context of UK agricultural AD characterised by high-solids feedstocks like maize silage and manure presents significant challenges. The standard ADM1 structure relies on Chemical Oxygen Demand (COD) as the universal unit for state variables due the original implementation was for wastewater AD systems [9–11]. While few studies have applied the COD-based framework to agricultural AD [12–14], doing so remains difficult in practice. For solid agricultural residues, the determination of COD balances is subject to substantial analytical uncertainty, since COD measurements require homogenisation and dilution of heterogeneous substrates, introducing significant experimental error. Unlike wastewater, the theoretical COD of complex composite substrates in agricultural AD is often only approximated, since the oxidation state of lignocellulosic biomass varies, thus creating stoichiometric uncertainty and producing COD gaps where substrate mass does not close the COD balance [9]. These challenges are further compounded by the common ADM1 assumption of constant liquid volume and density, which does not reflect the physical behaviour of high-solids reactors (TS > 10%), where degradation of solids into the gas phase results in measurable reductions in slurry mass and volume that influence hydraulic retention time (HRT) and washout rates of slow-growing methanogens [15]. To address these limitations, recent advancements have proposed mass-based modifications to the ADM1 framework, which align more closely with standard agricultural laboratory analyses and better represent the physical conservation of mass in solid AD [9,16–18].

### 1.3 Methodological Framework: Using Laboratory-Scale Proxies for Model Calibration

Parallel to these modelling advancements is the rise of the digital twin concept within the Industry 4.0 [19]. A digital twin is widely defined as a virtual replica of a physical system that is continuously updated with real-time data to mirror the condition and behaviour of the physical system throughout its entire operational life [20]. However, the usefulness of a digital twin is directly linked to the fidelity of its underlying process model. Before a model can serve as a reliable digital replica for a full-scale industrial plant, it must be rigorously calibrated and validated [21].

This study introduces the concept of the ‘physical twin’. A physical twin is not merely the industrial site itself, but can also refer to a highly relevant, scaled-down laboratory reactor that replicates the operational conditions of the full-scale site with high fidelity [21]. By conducting experiments on a physical twin, high-quality and frequency data can be generated under controlled conditions that are infeasible to achieve at the full industrial scale. The physical twin, once validated to sufficiently represent the full-scale system, can be used to generate data in regions with little or no data to validate modelling and control approaches. The concept is highly linked to multi-fidelity experimental and simulation studies. Multi-fidelity frameworks have traditionally been restricted to offline experimental design, such as asynchronous batch Bayesian optimisation for battery electrolytes [23]. This study presents a novel application of these frameworks by leveraging the physical twin explicitly for real-time process control and dynamic model validation. This data becomes a foundation for calibrating complex models such the ADM1, which can then be applied to improve efficiency at the full-scale facility. The use of these systems reduces overfitting and allows for more confidence for operating AD sites in new operating regimes. A schematic illustrating physical and digital twin as an integrated platform is shown in Figure 1.

**Figure 1.**
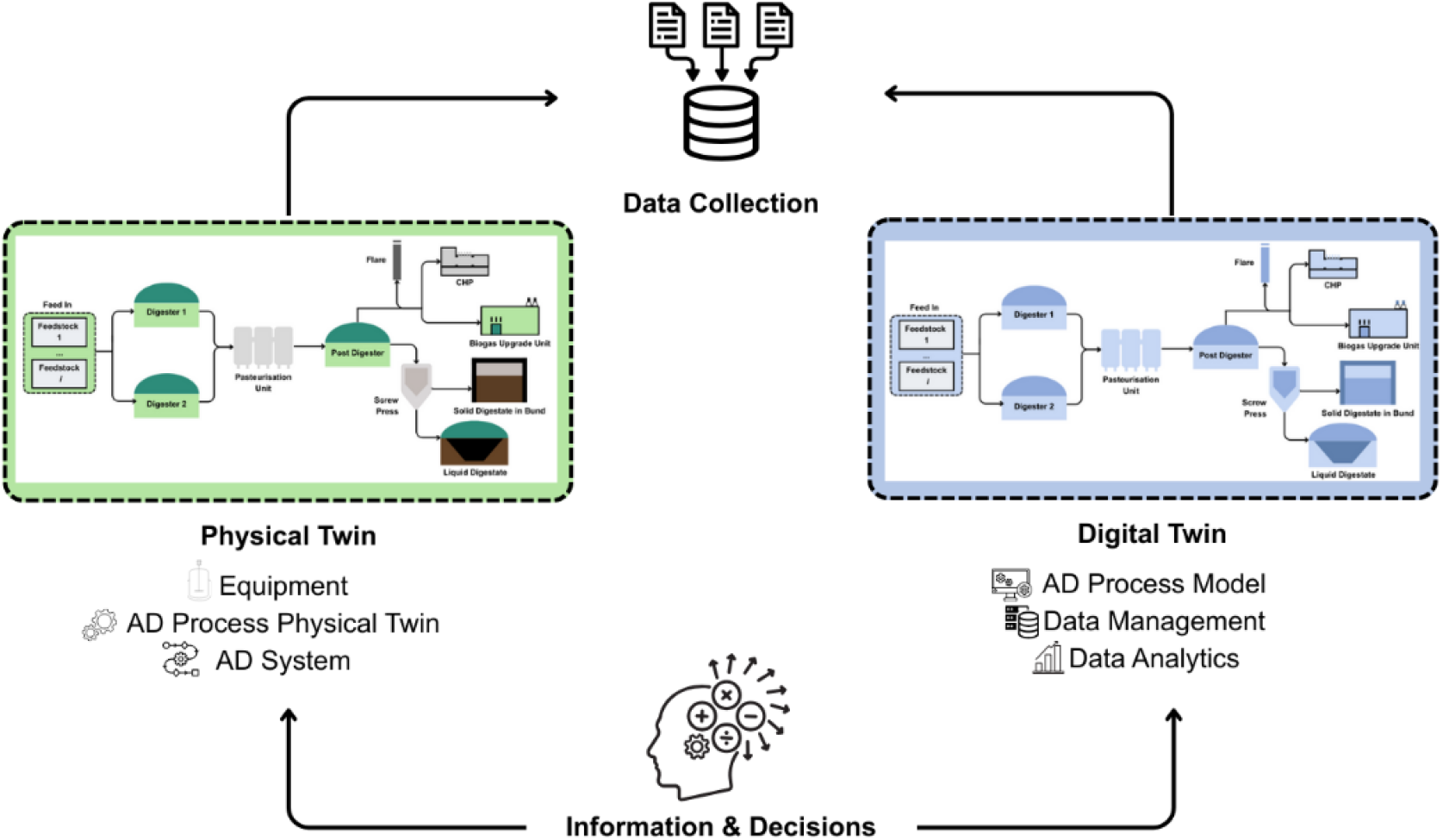
Schematic of integration of physical twin and digital twin adapted of AD system developed based of [22].

### 1.4 Research Novelty and Objectives

This study presents the employment and validation of a mass-based ADM1 model for a laboratory physical twin of a UK agricultural AD site co-digesting maize silage and cow manure. Most existing studies calibrate and validate models using data from a single reactor, often by dividing one continuous dataset into temporal segments. This approach increases the risk of overfitting, as the calibrated parameters may simply memorise reactor-specific noise and unmeasured disturbances rather than capturing the underlying system dynamics [24]. In this study, we use two identical 5-litre laboratory reactors acting as small-scale physical twins of the full-scale site. The first reactor is used for calibration under steady-state and dynamic conditions. The second reactor is maintained as an independent replica and is subjected to a distinct process disturbance by adjusting the OLR. The model, calibrated solely on the first reactor, is then used to predict the dynamic response of the second reactor. By validating the model against an independent twin subjected to different loading stresses, this approach tests the structural validity of the model and its ability to generalise to new operating conditions which is rarely achieved in literature, to the best of the authors knowledge. Furthermore, the ADM1 model is high-dimensional, therefore we integrated Global Sensitivity Analysis (GSA) using the Sobol method to systematically identify critical kinetic parameters, ensuring that the calibration process is focused and physically meaningful. By combining a mass-based model structure, a physical twin experimental design, and a cross-reactor validation approach, this study establishes a comprehensive methodological foundation for developing reliable predictive tools for the UK biogas sector.

The remainder of this paper is structured as follows: Section 2 provides details of the methodology, specifically describing the physical configuration and operational parameters of the bioreactor alongside the mathematical structure of the model. The core findings are presented and analysed in Section 3, which covers global sensitivity analysis, parameter estimation, and a comparative discussion of simulation results for the laboratory physical twins R6 and R5. Finally, Section 4 concludes the paper by summarising key insights and the implications of this study.

## 2.0 Methodology

### 2.1 Laboratory Physical Twin Bioreactor Configuration and Operation

Experimental data were obtained from a laboratory-scale semi-continuous AD physical twins with a total volume of 6 L and working volume of 4 L [25]. Two sets of reactors were operated in parallel to simulate full-scale biogas plant operation, each fed with a combined substrate containing 29 to 31 % total solids. The reactors were operated under mesophilic conditions ranging from 35 to 55 °C and continuously mixed with an asymmetric anchor-blade stirrer at 40 rpm to ensure homogeneous substrate distribution and prevent sedimentation.

The digester was inoculated with anaerobic sludge obtained from a full-scale agricultural biogas plant, providing an established microbial consortium adapted to lignocellulosic feedstocks. Feeding was performed every two days in semi-continuous mode, with a fixed volume of digestate removed and replaced with fresh substrate to maintain a constant HRT [25]. Figure 2 shows the physical twin experimental set-up.

**Figure 2.**
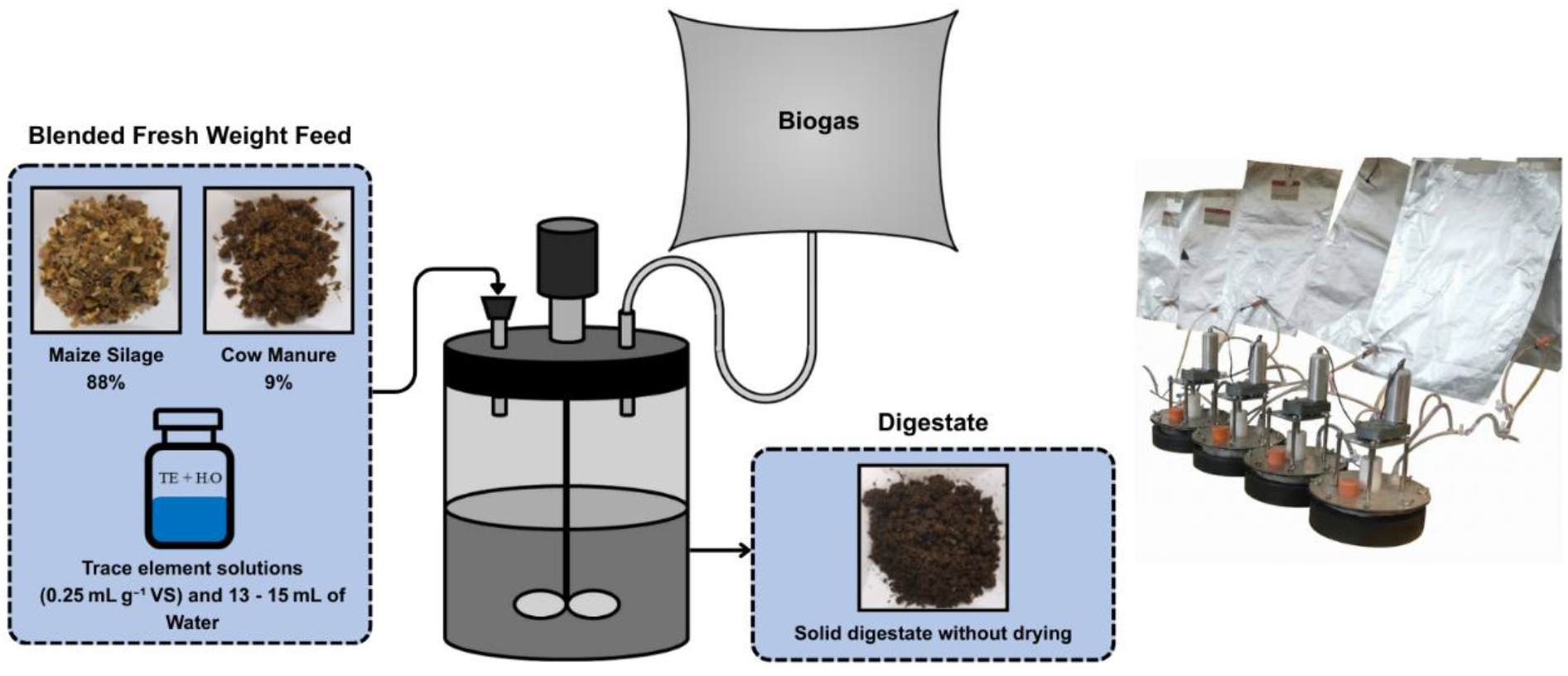
Laboratory Physical Twin experimental set-up of a full scale UK AD plant. A 6 L reactor feeding maize silage and cow manure with dried weight composition of 93 % and 7 % respectively.

The reactor was fed with a co-digestion mixture consisting primarily of maize silage with cow manure as co-substrate. On a fresh weight basis, the feed composition was maize silage (88%), cow manure (9%), and water and trace elements (3%). Trace element solutions (0.25 mL g^−1^ VS) and 13 to 15 mL of water were added every two days to compensate for evaporation. Crop silages were pre-processed to a particle size < 10 mm and stored at –20 °C until use [25]. Prior to day 202 of operation, the dried weight composition was maize silage (93 %) and cow manure (7 %), which was maintained after a feedstock adjustment on day 202, where the total solids content of maize silage increased from 32 % to 34 %, while cow manure content remained constant at 23 %. This feedstock composition was selected to mimic the operational conditions and substrate characteristics of a UK agricultural AD plant, allowing results from the laboratory physical twin reactors to represent full-scale digesters [25]. The OLR ranged from 3 to 4 g VS L^−1^ d^−1^, corresponding to total organic loads of 12 to 16 g VS d^−1^ and a solid retention time (SRT) of 75 to 115 days in the full-scale biogas plant. However, the OLR was also tested outside the usual range of the site scaled down to test the stress load on both physical twin reactors which is shown in Figure 3.

**Figure 3.**
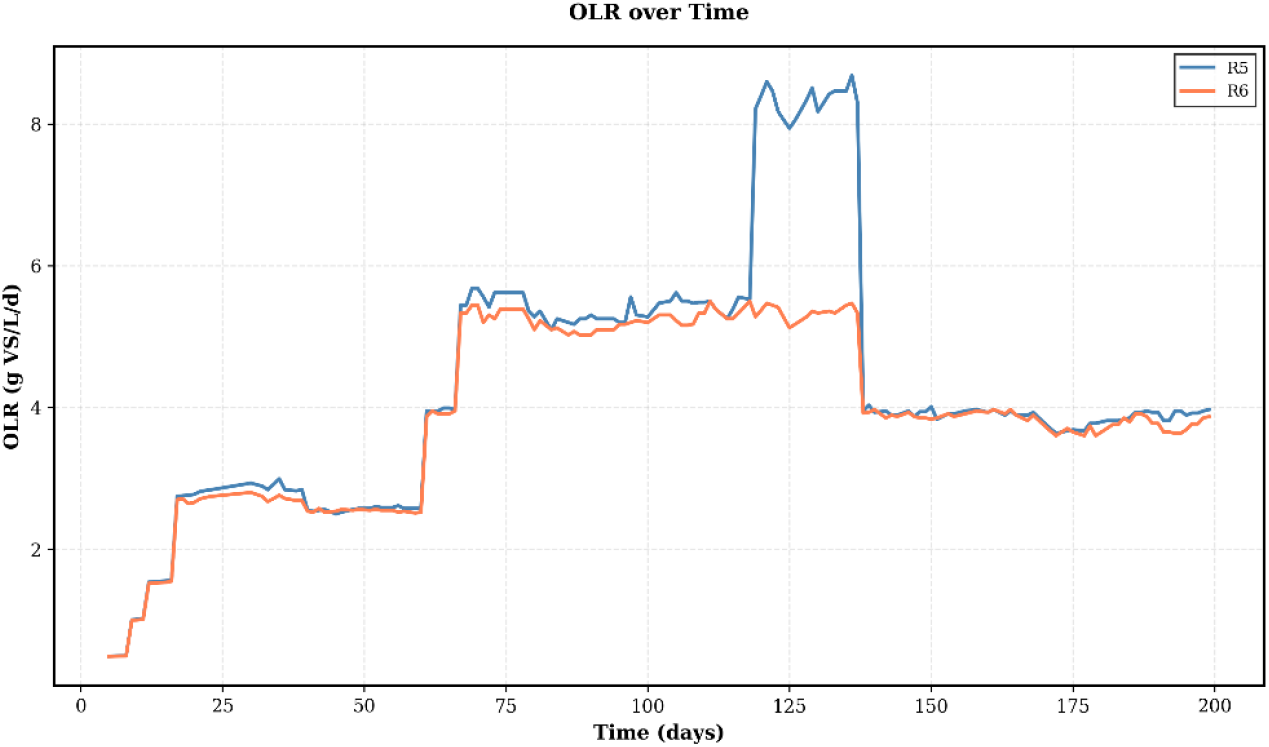
OLR over time of the physical twin reactors, the blue line is the R5 reactor with peak OLR reaching to 8.6 g VS L^−1^ d^−1^ and the orange line is the R6 reactor with peak OLR of 5.5 g VS L^−1^ d^−1^.

Substrate characterisation was performed during SRT and temperature changes to determine total solids, volatile solids, ash content, total and soluble chemical oxygen demand, total Kjeldahl nitrogen, total ammonia nitrogen, pH, and elemental composition (carbon, hydrogen, oxygen, nitrogen, sulphur, phosphorus). Additionally, organic polymer fractions including total proteins, starch, crude lipids, cellulose, hemicellulose, and lignin were quantified. VFA content and biochemical methane potential were also measured. All analyses were performed according to standard methods with thorough details presented in [25]. The OLR, SRT and temperature were systematically varied throughout the experimental period to assess reactor performance under different operational conditions and to generate data across a range of steady states.

The reactor was monitored continuously over the experimental period with daily measurement of biogas production volume using a gas flow meter. Biogas composition, specifically methane and carbon dioxide content, was analysed three times weekly by gas chromatography to track methanogenic activity and process stability. Liquid phase samples were collected twice weekly from the reactor effluent. These samples were immediately centrifuged and filtered for analysis of pH, VFA concentrations (acetate, propionate, butyrate, valerate), ammonia nitrogen, and alkalinity. VFAs were quantified by high-performance liquid chromatography to monitor the balance between acidogenesis and methanogenesis. The total and volatile solids concentrations in the reactor were measured weekly to track biomass accumulation and substrate degradation dynamics. The overall feedstock composition and characteristics measured are outlined in **Table 1**.

**Table 1:** Feedstock Characteristics of Maize Silage, Cow Manure and Mixed Co-digested Feedstock Blend.

| Feedstocks | Maize Silage | Cow Manure | Mixed |
| --- | --- | --- | --- |
| pH | 3.643 | 8.633 | 3.983 |
| Carbohydrates (g/kg FW) | 279.2 | 179.5 | 272.2 |
| Proteins (g/kg FW) | 22.59 | 24.22 | 22.70 |
| Lipids (g/kg FW) | 7.682 | 3.465 | 7.387 |
| TS (%) | 31.81 | 22.91 | 30.05 |
| VS (%) | 30.65 | 20.48 | 28.82 |
| ash (%) | 1.156 | 2.430 | 1.236 |
| TCOD (g/kg FW) | 369 | 274.6 | 349.4 |
| sCOD (g/kg FW) | 35.94 | 10.31 | 32.55 |
| ThCOD (g/kg FW) | 432 | 322 | 409.1 |
| EBMP (NmL/g VS) | 334 | 181 | 324 |
| total VFAs | 1.844 | 2.134 | 1.815 |
| acetic acid | 1.836 | 1.738 | 1.772 |
| propionic acid | 0 | 0.3634 | 0.0327 |
| i-butyric acid | 0 | 0 | 0 |
| n-butyric acid | 0 | 0 | 0 |
| i-valeric acid | 0.007935 | 0.03212 | 0.009873 |
| n-valeric acid | 0 | 0 | 0 |
| TKN (g/kg FW) | 4.001 | 4.697 | 3.943 |
| TAN (g/kg FW) | 0.3871 | 0.8223 | 0.4147 |
| total proteins (%TS) | 7.101 | 10.57 | 7.343 |
| total starch (%TS) | 25.66 | 0.2173 | 23.88 |
| crude lipids (%TS) | 1.548 | 0.1166 | 1.665 |
| cellulose (%TS) | 13.85 | 1.043 | 14.9 |
| hemi-cellulose (%TS) | 17.08 | 1.286 | 18.37 |
| lignin (%TS) | 13.89 | 1.045 | 14.94 |
| N free extract (%TS) | 39.76 | 2.993 | 42.76 |
| Carbon (%) | 44.9 | 43.15 | 44.78 |
| Hydrogen (%) | 6.55 | 5.72 | 6.492 |
| oxygen (%) | 43.38 | 38.32 | 43.03 |
| Nitrogen (%) | 1.15 | 1.83 | 1.198 |
| sulphur (%) | 0.07 | 0.37 | 0.091 |
| C/N | 39.11 | 23.7 | 38.03 |
| TP (g P/kg TS) | 2.432 | 1.7 | 2.381 |
| ortho-phosphate, PO<sub>4</sub>-P (g P/kg FW) | 0.5157 | 0.04935 | 0.4583 |
| ortho-P (g P/ kg TS) | 1.622 | 0.2154 | 1.525 |

The feedstock characteristics of the mixed feedstock in **Table 1** was calculated from the individual feedstocks based on the ratio of the individual feedstock contributing to the mixed feedstock.

### 2.2 Mathematical Model Structure

#### 2.2.1 Anaerobic Digestion Model No.1 (ADM1)

Since agricultural biomass and waste serve as the study’s feedstock, we chose to employ the mass-based ADM1 implementation developed by [9] which has proven to be suitable for agricultural AD over the COD based ADM1 developed for wastewater systems. The model is built upon a system of differential equations that track the concentration of 34 state variables. It separates the physicochemical processes (ion dissociation, gas-liquid transfer) from the biochemical reactions. The biochemical matrix uses Monod-type kinetics to describe the growth and decay of biomass groups, modulated by pH, temperature, and inhibition functions [26]. The four successive steps of AD which include: hydrolysis, acidogenesis, acetogenesis, and methanogenesis are described by the 34 state variables that make up the standard ADM1 structure [27]. The model has seven functional groups which are: sugar, amino acid, long-chain fatty acid, valerate and butyrate, propionate, acetoclastic methanogenesis, and hydrogenotrophic methanogenesis. These are used in the model to represent microbial activity in which each of these groups is responsible for transforming a particular substrate [27,28]. Figure 4 represents model implementation of the inputs and outputs of the ADM1.

**Figure 4.**
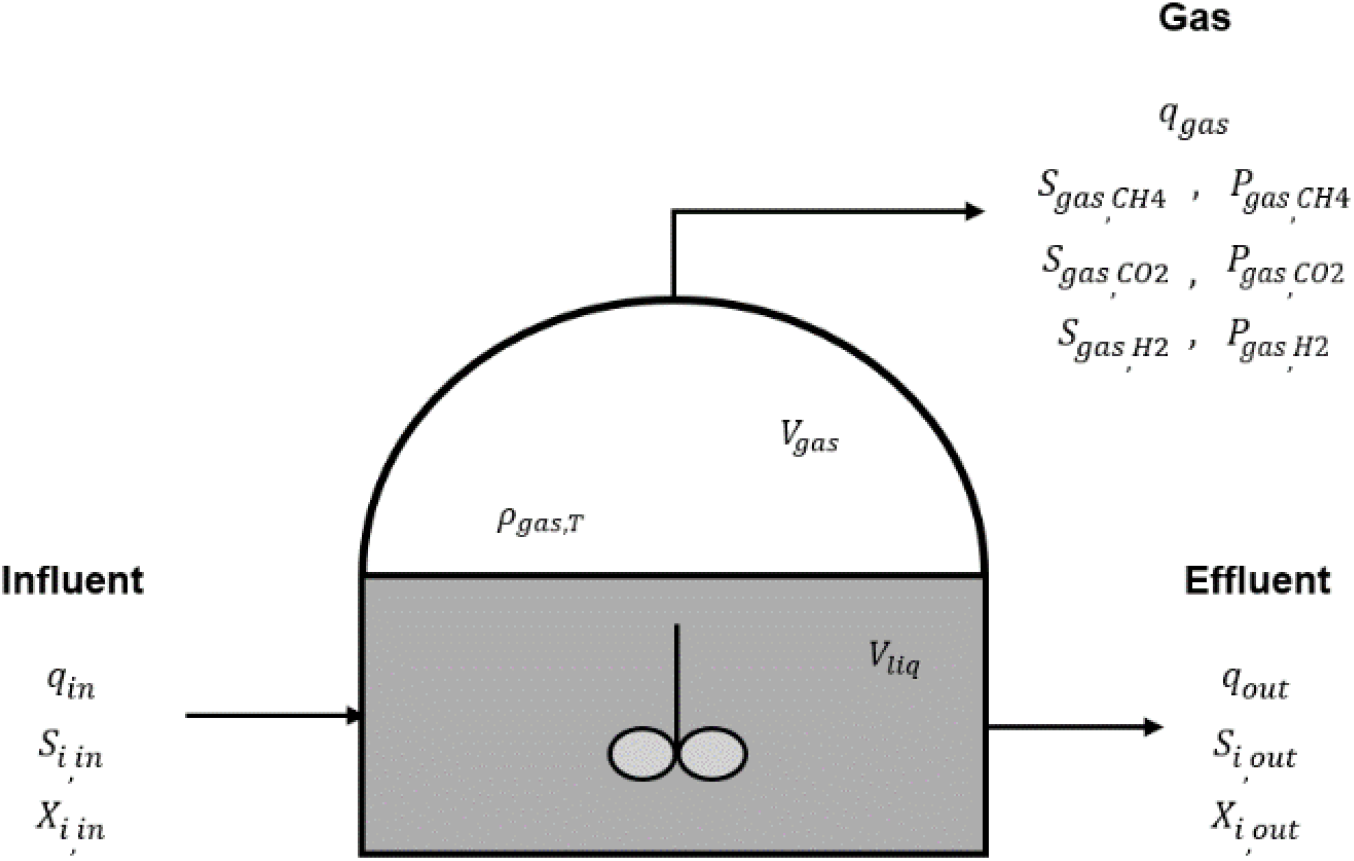
Schematic of Continuous Stirred Tank Reactor (CSTR) implementation of the ADM1 model based of [6].

The input and output flows for the ADM1 in Figure 4 represent ***S***_***i***,***in***/***out***_= soluble components concentration in the input and output, ***X***_***i***,***in***/***out***_= particulate components concentration in the input and output, ***q***_***in***_, ***q***_***out***_, ***q***_***gas***_ are flow rates in, out and gas flowrate, **S**_**gas**,**CH4**_, **S**_**gas**,**CO2**_, **S**_**gas**,**H2**_ are concentration of individual gas components, 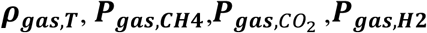 are total pressure and pressure of individual gas components, ***V***_***liq***_, ***V***_***gas***_ are volumes in liquid and gas phase.

A simplified derivation of the mass based ADM1 model is outlined below stating the components and equations involved adapted from [29]. A more thorough and detailed construction of the model equations can be found in [9]. By applying the principle of mass conservation to a CSTR, the equations governing the system can be expressed as follows.

##### 2.2.1.1 Liquid and Gas Phase Mass Balances

The accumulation of soluble components (*S*_*i*_) in the reactor’s liquid volume (*V*_*liq*_) is determined by the influent and effluent flow rate (*F*), liquid concentrations (*S*_*i*_), and internal reaction rates (*ρ*_*j*_). As shown in Eq.1, this balance accounts for soluble species including sugars (*S*_*su*_), amino acids (*S*_*aa*_), fatty acids (*S*_*fa*_ to *S*_*ac*_), and dissolved gases 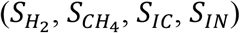:

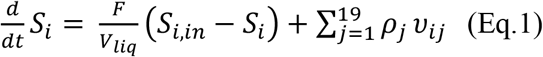

Similarly, the balance for particulate components (*X*_*i*_ ) is described in Eq.2. These include composite substrates (*X*_*ch*_, *X*_*pr*_, *X*_*li*_) and microbial biomass groups (*X*_*su*_ to 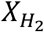):

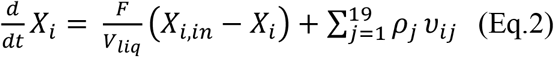

For the gas phase, the concentration of gas components (*S*_*gas,i*_), specifically *H*_2_, *CH*_4_, and *CO*_2_, is governed by the flow rate and the specific mass transfer rate (*ρ*_*T,i*_) between the liquid and gas volume (*V*_*gas*_) as shown in Eq.3:

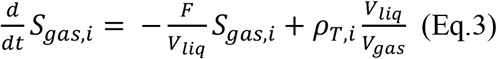

The specific mass transfer rate described in Eq.4 is determined by the mass transfer coefficient (*K*_*La*_), the Henry’s law coefficient (*K*_*H*_), and the liquid concentrations (*S*_*liq,i*_):

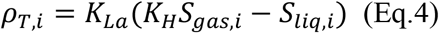

##### 2.2.1.2 Gas Dynamics and Headspace Pressure

The partial pressure of pressures for methane 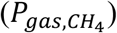 in Eq.5, carbon dioxide 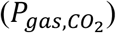 in Eq.6, and hydrogen 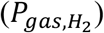 in Eq.7 components are calculated using the ideal gas law at the system temperature (*T*_*ad*_) and the universal gas constant (*R*):

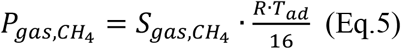

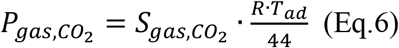

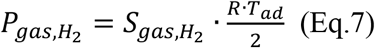

Assuming that water vapor has saturated the reactor headspace at a constant partial pressure (*P*_*gas,H*2*O*_), the total headspace pressure (*P*_*gas*_) is calculated as the sum of the partial pressures of *CH*_4_, *CO*_2_, *H*_2_, and *H*_2_*O*. The gas production rate *q*_*gas*_ shown in Eq.8 is then determined by the pressure difference between the reactor headspace and the atmospheric pressure (*P*_*atm*_), governed by the gas flow resistance constant (*k*_*p*_):

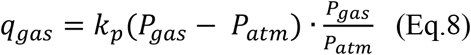

The methane flow rate (*q*_*ch*4_) shown in Eq.9 is then deduced as a fraction of the total flow based on its partial pressure:

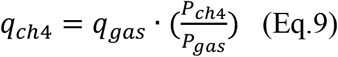

##### 2.2.1.3 Ionic Components and Biochemical Kinetics

Ion components, including cations (*S*_*cat*+_) and anions (*S*_*an*−_), are tracked via Eq.10 and Eq.11, while Eq.12 describes the reaction rates for dissociated species (*S*_*i*−_) which represents the concentrations of 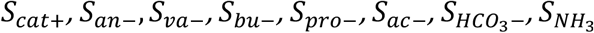.

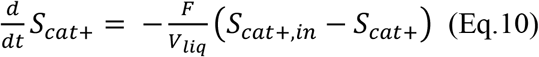

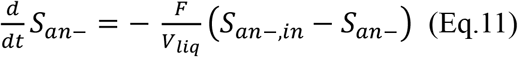

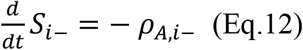

The ADM1 framework categorises 28 processes into 19 biochemical processes and 9 physical processes. Biomass growth is modelled using Monod-type functions (Eq.13) incorporating the maximum uptake rate constant (*k*_*m,i*_) and half-saturation constant (*K*_*S,i*_), while decay follows first-order kinetics (Eq.14) with a decomposition rate constant (*k*):

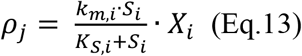

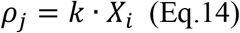

The pH value (Eq.15) is calculated based on the hydrogen ion concentration (*S*_*H*+_) as follows based on Eq.16 and Eq.17:

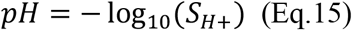

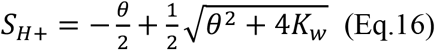

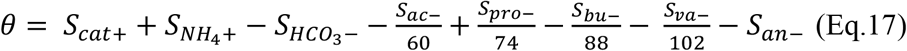

Where the charge balance (*θ*) incorporates cations, ammonia 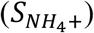, bicarbonate 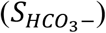, anions, and the various VFA fractions, and *K*_*w*_ which represents the hydrogen ion transfer rate.

##### 2.2.1.4 Inhibition Framework and Operational Stability

The model incorporates inhibition coefficients (*I*) to account for environmental stressors. pH inhibition (*I*_*pH*_) is defined by lower (*pH*_*LL*_) and upper (*pH*_*UL*_) limits as shown in in Eq.18 – Eq.20:

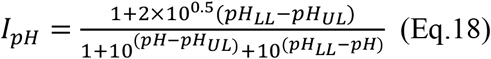

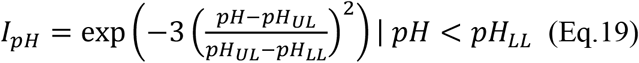

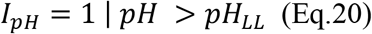

Finally, secondary inhibitions include valerate/butyrate competition (*I*_1_) is described in Eq.21, limited inorganic nitrogen (*I*_*IN,lim*_ ) in Eq.22, dissolved hydrogen (*I*_*H*2_ ) in Eq.23, and free ammonia 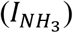 in Eq.24:

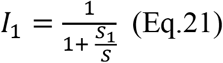

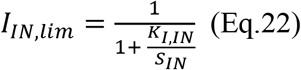

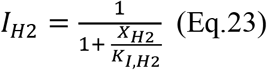

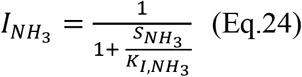

To accurately simulate the AD process using the ADM1 model, it is essential to first determine the key parameters that significantly affect its performance, such as the influent concentrations (*S*_*i,IN*_, *X*_*i,IN*_) and certain rate constants. Additionally, inhibition factors such as *K*_*I,IN*_, *K*_*I,H*2_, and *K*_*I,NH*3_ must be identified, especially when considering a biogas plant feed with specific types of organic material. Once these parameters are identified and adjusted, the model can be reliably used to simulate the AD process.

Simulations have demonstrated that the time constants for soluble, particulate, and gas components are substantially longer than those for ion components [29]. Consequently, the soluble, particulate, and gas components might be considered as slow-changing, whereas the ion components are fast-changing. Additionally, certain slow-changing components are not extremely sensitive to changes in the substrate supply, indicating that variations in the feed have limited influence on these components which is useful when optimising the AD process, of the key components that require attention for maximum performance [29].

To evaluate the operational stability and performance of the anaerobic digestion system, several algebraic equations were integrated into the ADM1 framework based on [6,30,31]. These outputs provide a bridge between the model’s state variables and common laboratory measurements.

##### 2.2.1.5 Alkalinity and Buffer Capacity

To evaluate the operational stability and performance of the anaerobic digestion system, several algebraic equations were integrated into the ADM1 framework based on [6,30,31]. These outputs provide a bridge between the model’s state variables and common laboratory measurements. The alkalinity components are crucial for determining the reactor’s resistance to pH acidification.

The Partial Alkalinity (*PA*) represents the buffering capacity provided by the bicarbonate and ammonia systems, defined by the sum of the inorganic carbon and nitrogen states as shown in Eq.25:

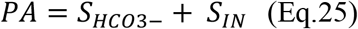

The Intermediate Alkalinity (*IA*) : represents the buffering capacity associated with VFAs described in Eq.26. This calculation accounts for both dissociated and undissociated forms to align with standard titration methods. The division by molecular weights (60, 74, 88, 102) converts the mass concentration (g/L) to molar concentration (mmol/L), while the factor of 1000 to express the result in g *CaCO*_3_-equivalent/L.

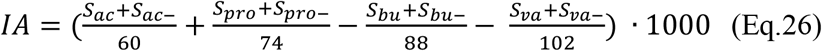

Finally, the Total Alkalinity ( *TA* ) is calculated via the system’s charge balance (Eq.27), incorporating inorganic nitrogen, bicarbonate, dissociated VFAs, and the balance of anions and cations:

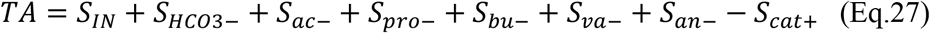

Together, these calculated alkalinity fractions allow for the continuous evaluation of the reactor’s chemical buffering state, providing a vital metric for the early detection of kinetic imbalances and process instability.

##### 2.2.1.5 Organic Load and Effluent Quality

The following equations characterise the organic concentrations within the reactor. Chemical Oxygen Demand (COD) was purely for the physical twin results comparison despite using the mass based ADM1 via conversion factors. The conversion factors suggested by [32] are outlined in **Table 2**.

**Table 2:** State conversion factors suggested by [32] used for VFA and COD determination for physical twin results comparisons.

| State Conversion Factors | Value (g COD/g) |
| --- | --- |
| $COD_{ac}$ | 1.0667 |
| $COD_{pro}$ | 1.5135 |
| $COD_{bu}$ | 1.8182 |
| $COD_{va}$ | 2.0392 |
| $COD_{su}$ | 1.0667 |
| $COD_{aa}$ | 1.42 |
| $COD_{fa}$ | 2.875 |
| $COD_{ch}$ | 1.1841 |
| $COD_{pr}$ | 1.5304 |
| $COD_{li}$ | 2.8609 |

The total mass concentration of VFAs, expressed in COD terms (Eq.28), is determined by the sum of acetate, propionate, butyrate, and valerate concentrations multiplied by their respective conversion factors:

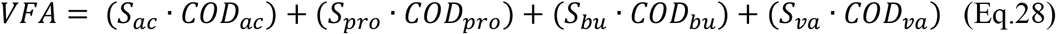

Total COD represents the sum of all organic components within the liquid phase as shown in Eq.29. The model accounts for the conversion of particulates into soluble forms through hydrolysis, which contributes to the final liquid-phase COD concentration:

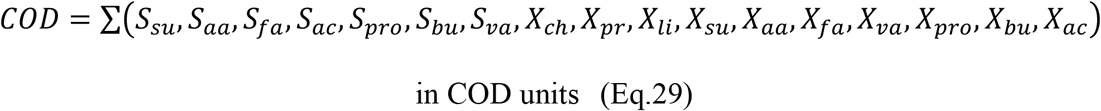

Finally, Eq.30 describes the Total Ammonia Nitrogen (*TAN*) is directly mapped from the inorganic nitrogen state variable:

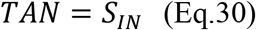

However, MATLAB was used to construct the mass-based ADM1. With the aid of the openly accessible Python implementation of the original ADM1 [33] model, known as “PyADM1,” the MATLAB implementation was converted to Python using the SciPy library. Using Python’s scientific computing modules for data analysis and numerical integration, this translation preserved all mathematical formulations and parameter values from the mass-based model.

To resolve the system of 34 ordinary differential equations (ODEs), the scipy.integrate.solve_ivp function was used, employing the Backward Differentiation Formula (BDF) method. This implicit solver was specifically chosen to handle the numerical stiffness inherent in ADM1’s rapid biochemical and acid-base transitions. The simulation maintained high numerical precision with a relative tolerance (rtol) of 10^-6^ and an absolute tolerance (atol) of 10^-8^. The simulation time steps were discretised according to the frequency of the influent data, which was daily, while the solver used internal adaptive stepping to ensure convergence within each interval.

##### 2.2.1.6 ADM1 Model Input

To use the ADM1 model, the feedstock characteristics were used to define the ADM1 model inputs over time. The list of the values of the most important ADM1 states used were determined from the approach developed by [34] are outlined in **Table 3**.

**Table 3:**
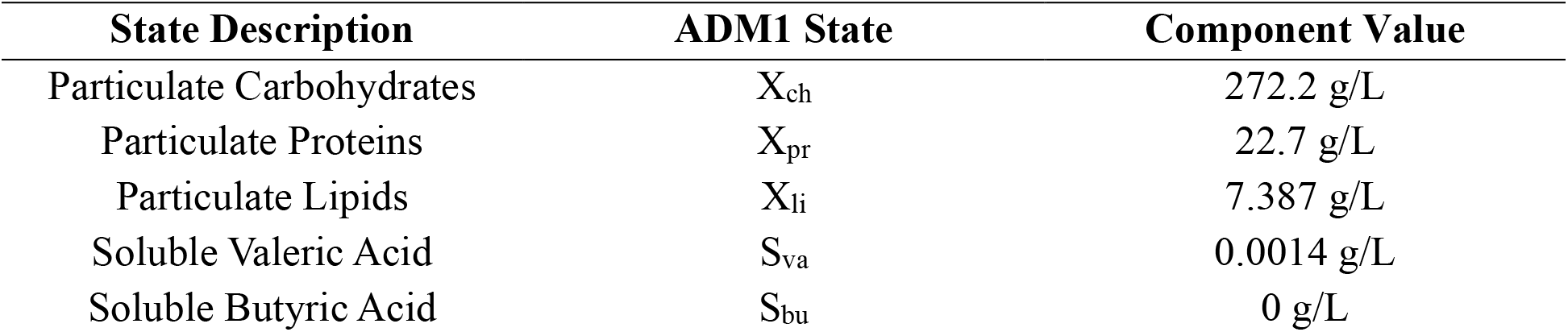
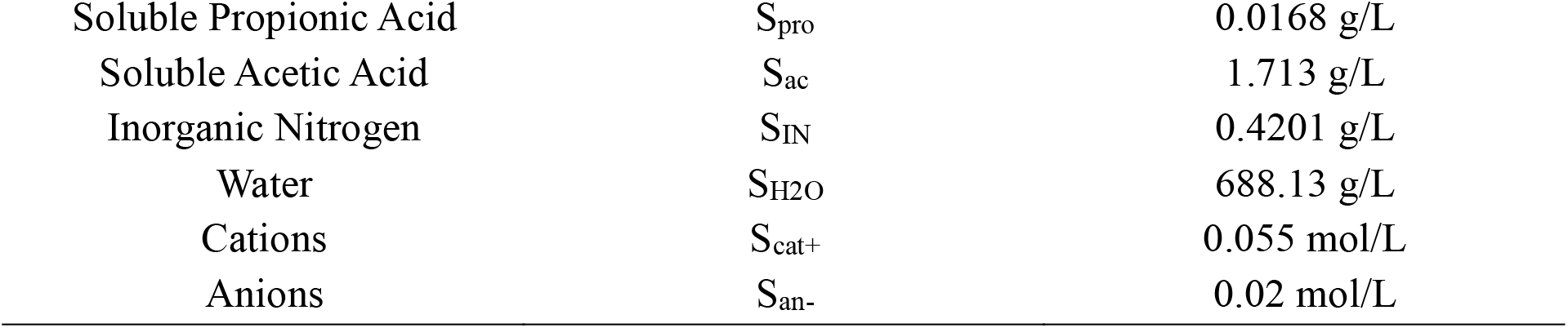
List of states within the ADM1 model framework.

The following states were the minimum states required to run the model suggested and validated by [34]. The values for S_cat+_ and S_an-_ were manually adjusted as suggested from [17].

#### 2.2.2 Parameters and Calibration

This section details the hierarchical framework employed for model calibration. To address the challenge of over-parameterisation, the process begins with a Global Sensitivity Analysis (GSA) to identify the subset of parameters with the most significant impact on model output [35]. Subsequently, the parameter estimation problem is formulated to define the goodness-of-fit metrics, followed by the description of the specific optimisation procedure used to minimise the error between simulated and experimental data.

##### 2.2.2.1 Global Sensitivity Analysis

GSA using the variance-based Sobol method [36] was used to identify the most influential kinetic parameters with respect the mass based ADM1 structure. While GSA has been applied previously to standard COD-based ADM1 formulations by [37,38], to the authors’ knowledge, it has not been conducted for full ADM1 mass-based implementation. Although previous research by [9], proposed simplified versions of the mass-based model, this study deliberately retains the full structural complexity. This approach is necessary to simultaneously capture and predict multiple process outputs monitored within the physical twin. We wanted to use the model which can provide the maximum outputs from the process. This step is essential because the shift from a COD basis to a mass basis alters stoichiometric relationships and reaction rates. As a result, parameters that are influential in COD-based models may not retain the same significance in a mass-based formulation. Conducting GSA under the updated modelling framework therefore ensures that calibration efforts remain focused on parameters that that drive variance in the mass-based outputs and are aligned with the structural changes introduced in the model.

The Sobol method is a model-independent, variance-based approach that decomposes the variance of the model output into fractions which can be attributed to inputs or sets of inputs [36]. Considering the model as:

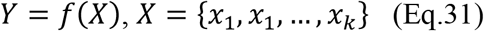

where *X* is a vector of *k* uncertain model parameters, the total variance of the model output *V*(*Y*) can be decomposed as:

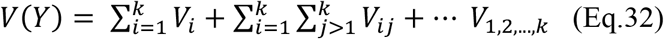

where *V*_*i*_ represents the variance contribution of the parameter *x*_*i*_ alone, and subsequent terms capture the variance contributions due to interactions between parameters.

Two sensitivity indices are calculated for each parameter. The first-order sensitivity index (*S*_*i*_): Measures the main effect of parameter *x*_*i*_ on the output variance without considering interactions with other parameters:

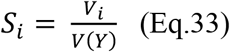

The total-order sensitivity index (*S*_*T,i*_) measures the total contribution of parameter *x*_*i*_, to the output variance, including both its direct effect and all interactions with other parameters:

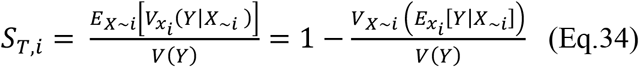

where *X*_∼*i*_ denotes the set of all parameters except *x*_*i*_.

Given the complexity of the ADM1 structure, the total-order index ( *S*_*T,i*_ ) is particularly valuable as it accounts for the non-linear coupling between biochemical processes and identifies parameters that may only become influential through their interaction with others.

The analysis was computationally implemented in Python using the open-source SALib library [39]. To ensure numerical stability and efficient convergence of the indices, the parameter space was sampled using the Saltelli sampling scheme [40], which extends the Sobol sequence to reduce error rates in the calculation of variance-based indices. The total cost of the analysis, defined as the number of model evaluations, follows the relation *N*_*total*_ = *N*(2*D* + 2), where *D* is the number of uncertain parameters and *N* is the base sample size [39]. For this study, only 25 kinetic and inhibition constants were selected out of the 60 parameters from the ADM1 mass-based model [9], as these represent the biologically sensitive variables governing the 19 biochemical processes (*D* = 25) . The remaining parameters which were fundamental physicochemical constants, equilibrium rates, numerical stability requirements, and design-specific parameters were kept fixed, as they do not vary with the microbial population or substrate type. A base sample size of parameters *N* = 512, was selected, resulting in a total of 26,624 simulations. This magnitude was chosen to ensure the confidence intervals of the sensitivity indices were sufficiently narrow to distinguish influential parameters from noise. A uniform probability distribution was used due to insufficient information describing the precise probability distribution of each parameter, as noted by [14].

##### 2.2.2.2 Parameter Estimation

Parameter estimation was performed by minimising the residual model deviation between the experimental data and the simulation results [41]. To automate this process and ensure efficient convergence within the parameter space, the Optuna hyperparameter optimisation framework which is a tree-structured Parzen Estimator with two probability models, [42] was used in Python.

The quality of the model fit was quantified using a single objective function, where Root Mean Square Error (RMSE) was selected in this study as the goodness-of-fit metric. The parameter estimation problem is formulated as an optimisation task where the objective is to minimise the objective function, *J*(*θ*), defined as:

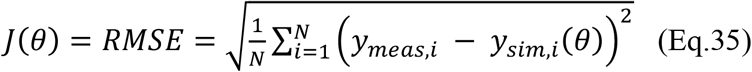

In this formulation, *N* represents the total number of experimental data points, while *y*_*meas,i*_ and *y*_*sim,i*_(*θ*) denote the experimental data and the corresponding model simulation values at point *i*, respectively. The vector of parameters to be identified, *θ*, is determined by the results of the GSA. By minimising *J*(*θ*), the calibration process ensures that the simulated outputs closely align with the observed dynamics of the physical system.

Based on the GSA results, exhibiting a total-order sensitivity index that cumulatively accounted for >95% of the output variance were considered for parameter estimation isolating the 10 most influential parameters across all modelled variables [35]. To ensure the model accurately reflects the experimental data, a systematic calibration approach was adopted which is illustrated in Figure 5.

**Figure 5.**
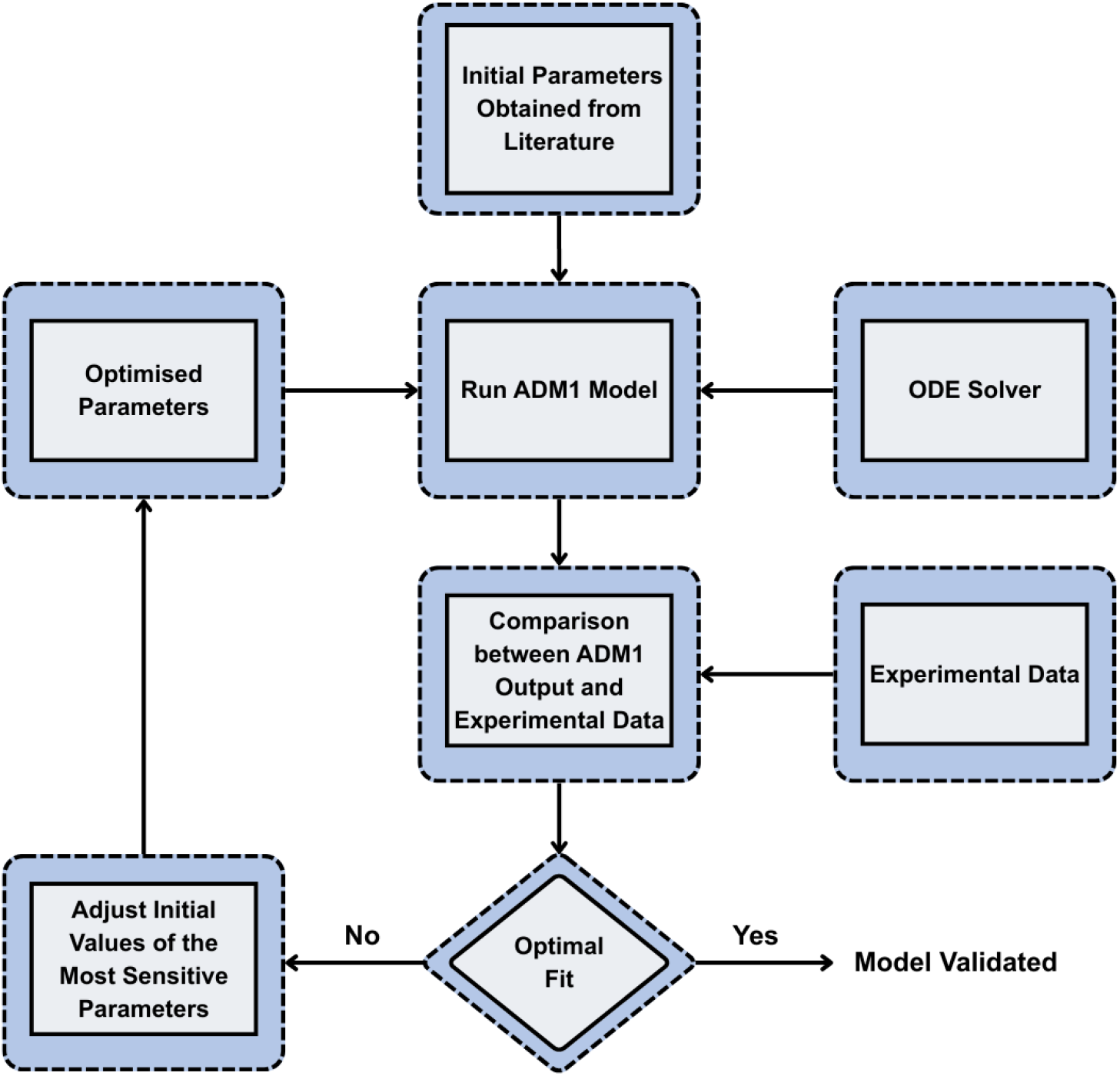
A schematic showing the model calibration of the ADM1 model using initial literature parameter values which are tuned to fit the measured the actual measurements.

In this workflow, Optuna acts as the external optimiser. For each iteration, the algorithm suggests a new set of values for the sensitive parameters (*θ*) based on the history of previous trials, aiming to minimise *J*(*θ*) . The solver evaluates the ADM1 model with these new parameters, calculates the RMSE against the experimental data, and returns this value to Optuna. This loop continues until the objective function converges to a minimum, providing the final optimal parameter set.

#### 2.2.3 Model Validation and Performance

First, the Root Mean Squared Error (RMSE) was employed to measure the average magnitude of the deviation between simulated and observed values as shown in Eq.36. This metric is particularly useful for identifying large outliers, as the errors are squared before being averaged:

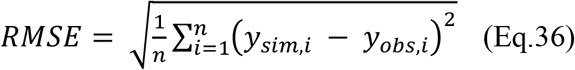

In this expression, *y*_*sim,i*_ represents the simulated value at time *i*, while *y*_*obs,i*_ denotes the corresponding observed (measured) value.

Second, the Mean Absolute Error (MAE) was used to provide a linear representation of the average error as shown in Eq.37. Unlike RMSE, MAE weights all individual differences equally, offering a straightforward assessment of the model’s average precision:

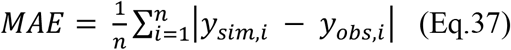

Finally, the Coefficient of Determination (*R*^2^) was calculated to evaluate the proportion of variance in the observed data that is successfully captured by the model as presented in Eq.38. This metric indicates the goodness-of-fit relative to the mean of the observed values 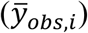:

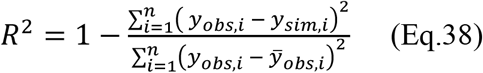

Together, these metrics provide a comprehensive statistical basis for assessing the model’s performance across both steady-state and transient operational phases. All simulations were performed on a workstation equipped with an AMD Ryzen 7 3700X processor (8 cores, 16 threads, 3.6 GHz), 32 GB RAM.

## 3.0 Results and Discussion

In this section the results of the GSA highlighting the top parameters which should be used for parameter estimation along with the optimal value for the parameters are shown. The simulation outputs follow for each physical twin reactor with performance metrics.

### 3.1 Global Sensitivity Analysis

The GSA aimed to quantify the influence of individual kinetic parameters on the mass-based ADM1 outputs and to prioritise them for parameter estimation. **Table 4** presents the sensitivity rankings derived from the total-order Sobol indices (*S*_*T,i*_) which considers both the direct effects of parameters and their interactions with other parameters.

**Table 4:**
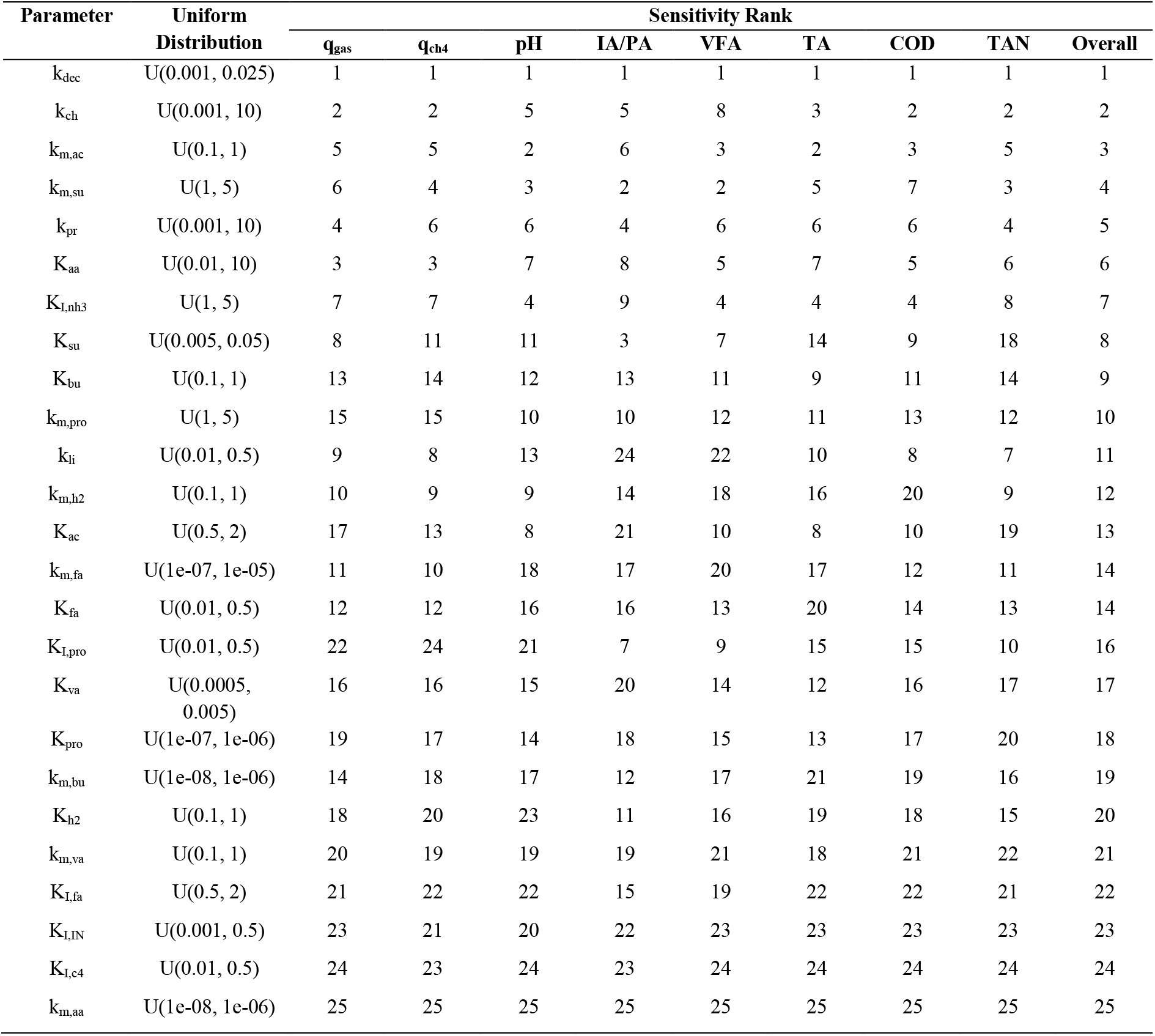
Parameter Rankings across all outputs (Sobol Total-Order *S*_*T,i*_). U(a,b) stands for the uniform distribution where a is the lower bound and b is the upper bound for the parameter.

The biomass decay rate (k_dec_) emerged as the most influential parameter, consistently holding the first rank across all individual outputs and the overall composite rank. This indicates that in a mass-based formulation, the variance in biomass maintenance and death is the primary factor governing the stability of the entire system, from gas production (q_gas_) to nutrient levels (TAN). While k_dec_ is globally dominant, substrate-specific kinetic rates show high sensitivity for relevant outputs. In terms of process kinetics, k_ch_ was identified as the second most influential parameter for both biogas production and COD, validating its role in governing the breakdown of complex solids. The model’s sensitivity to pH and VFA was mainly due to the acetate (k_m,ac_) and sugar (k_m,su_) uptake rates, which were ranked 2^nd^ and 3^rd^, where the conversion rates of these intermediate acids directly control the acid-base balance within the digester. In contrast, the negligible impact of inhibition constants (K_I,c4_) and amino acid kinetics (k_m,aa_), which consistently ranked at the bottom, indicates that the operating conditions did not push the system toward these specific performance-limiting thresholds. These findings align with GSA on the standard COD-based ADM1 models. Trucchia & Frunzo [43], focused on methane and VFAs, and a comprehensive review by Barahmand & Samarakoon [44], evaluated methane, COD, pH, and TAN across the literature identified disintegration/hydrolysis constants (k_dis_, k_ch_) and maximum specific uptake rates (k_m,ac_) as the key sensitive parameters. While this study similarly confirms the localised importance of k_ch_ and k_m,ac_ for gas and pH outputs, the global dominance of k_dec_ observed here highlights a fundamental characteristic of the mass-based formulation.

The results of the GSA provided a statistically robust basis for reducing the dimensionality of the subsequent model calibration. By identifying that a small subset of kinetic parameters accounts for the vast majority of output variance. The parameter estimation can be focused exclusively on these highly influential parameters while keeping non-influential parameters at their literature-standard values. This approach ensures that the model is not over-parameterised and that the resulting calibrated values are physically meaningful within the context of the mass-based ADM1 structure.

### 3.2 Parameter Estimation

Following the GSA, the highest-ranking kinetic parameters were calibrated by minimising the residual error between experimental observations and model simulations. The resulting calibrated values are presented in **Table 5**.

**Table 5:**
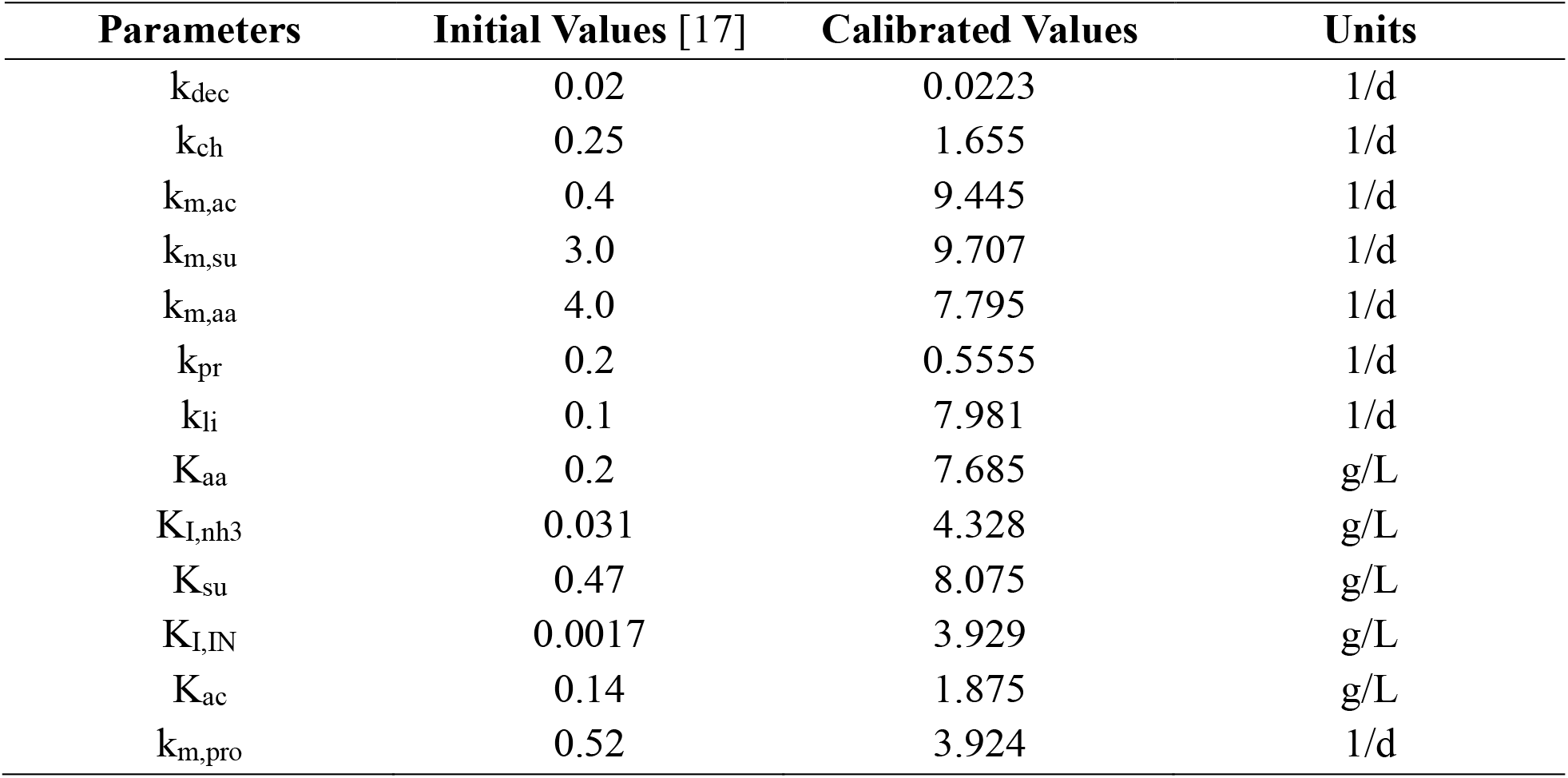
Initial Values used by [17], and calibrated values of top sensitivity ranked model parameters across all outputs.

These calibrated values provide a baseline for the mass-based ADM1, ensuring that the model accurately reflects the specific biochemical kinetics of the reactor while maintaining mass-balance across all state variables.

### 3.3 Simulation and Performance Results

#### 3.3.1 Laboratory Physical Twin R6

The performance of the calibrated mass-based ADM1 was evaluated by comparing the simulated outputs against experimental data obtained from Physical Twin R6 over a 160-day operational period with results presented in Figure 6. This duration was used to assess the model’s ability to predict dynamic shifts in reactor behaviour under varying OLRs and SRT conditions.

**Figure 6.**
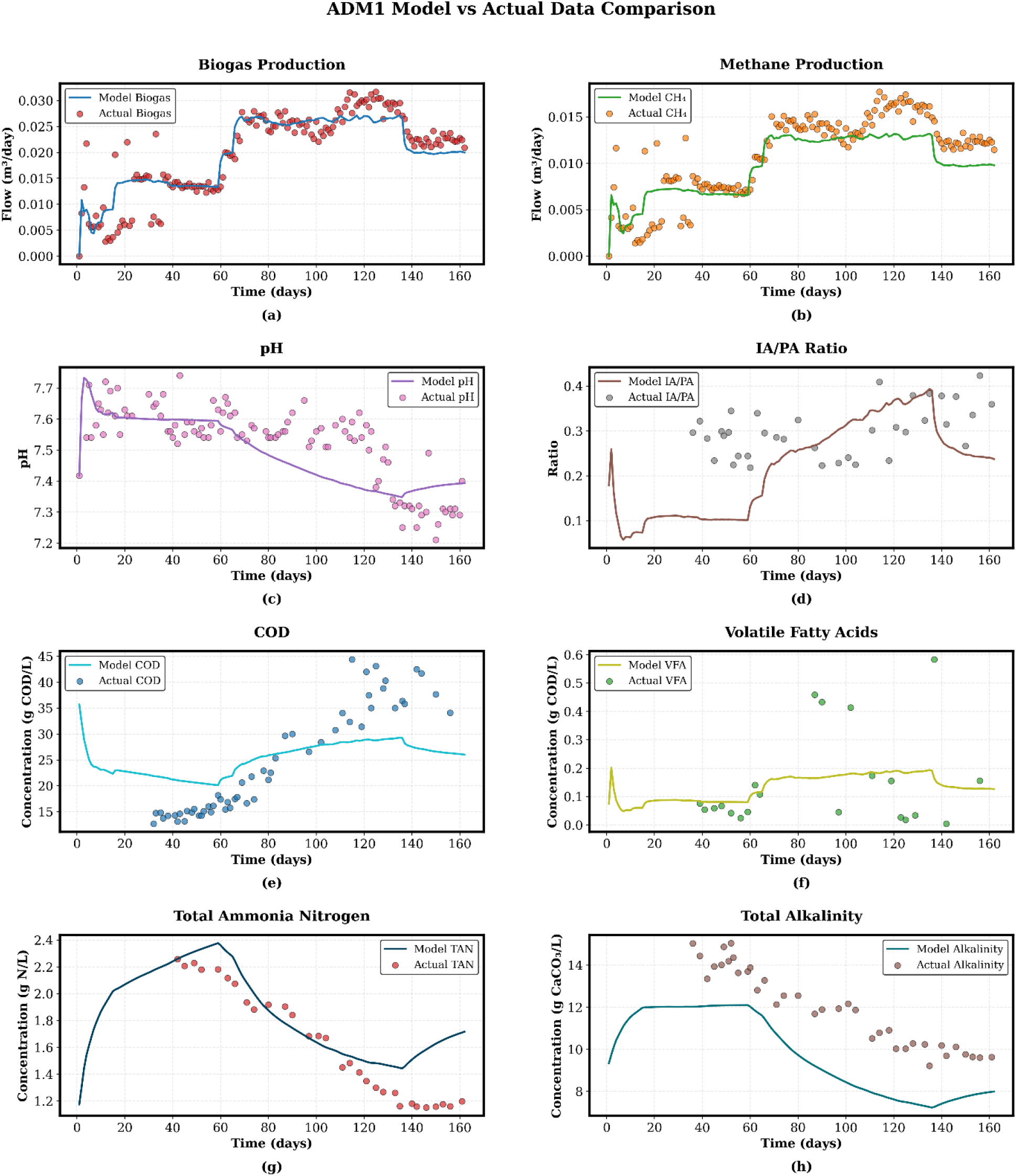
Simulation output and actual output comparison plots for key outputs of interest for physical twin reactor R6.

The model shows high fidelity in tracking both biogas q_gas_ and methane q_ch4_ production rates. It successfully captures the distinct step-changes in gas production occurring around day 60 and day 140, which correspond to changes in the influent feeding pattern. The close alignment between the model line and the experimental scatter points indicates that the calibrated kinetic parameters, particularly the carbohydrate hydrolysis rate k_ch_, accurately represent the rate-limiting steps of the AD process.

The simulated pH profile shows a steady decline from day 60 onwards, mirroring the experimental trend. While the model slightly overestimates the initial pH peak, it effectively captures the stabilisation around 7.4. Similarly, the TAN profile demonstrates the model’s capacity to track nutrient accumulation and release, peaking near day 60 before gradually declining in line with the actual data.

The COD simulations follow the baseline experimental values but show some deviation during the late-stage peak (days 120-140), where actual COD reached ∼45 g COD/L. VFA concentrations remained low throughout the period, which the model predicted accurately, confirming the presence of a well-balanced microbial community where intermediate production and consumption rates are similar.

To quantify the model’s predictive accuracy, statistical metrics including RMSE, MAE, and R^2^ were calculated for the key outputs in Table 6.

**Table 6:** Model Performance on the key model outputs for Physical Twin R6.

| Variable | RMSE | MAE | $R^2$ |
| --- | --- | --- | --- |
| $q_{\text{gas}}$ | 0.003407 | 0.00239 | 0.81 |
| $q_{\text{ch4}}$ | 0.002353 | 0.001962 | 0.71 |
| pH | 0.09814 | 0.07907 | 0.38 |
| TAC | 2.44 | 2.353 | -0.76 |
| IA/PA | 0.1233 | 0.1059 | -3.96 |
| COD | 7.33 | 6.447 | 0.48 |
| VFA | 0.1556 | 0.1077 | 0.13 |
| TAN | 0.2489 | 0.1969 | 0.61 |

**Table 7:** Model Performance on the key model outputs for Physical Twin R5.

| Variable | RMSE | MAE | R<sup>2</sup> |
| --- | --- | --- | --- |
| q<sub>gas</sub> | 0.0037 | 0.00259 | 0.85 |
| q<sub>ch4</sub> | 0.0028 | 0.0021 | 0.74 |
| pH | 0.1107 | 0.0939 | 0.62 |
| TAC | 2.193 | 1.959 | -0.69 |
| IA/PA | 0.2533 | 0.1672 | -9.07 |
| COD | 8.479 | 7.5 | 0.5 |
| VFA | 0.1762 | 0.1364 | -1.96 |
| TAN | 0.2816 | 0.2344 | 0.45 |

The model achieved high R^2^ values for gas production variables R^2^ > 0.81 for q_gas_ and 0.71 for q_ch4_, validating the models as a predictive tool for biogas and methane predictions. The lower R^2^ observed for pH with a value of 0.38 is largely attributed to the high sensitivity of the pH scale and the relatively small range of experimental variance. However, the low RMSE of 0.0981 confirms that the model tracks the overall pH trends well within a highly acceptable range for monitoring reactor operation. The negative R^2^ values for TAC and IA/PA suggests that while the trends are captured, the high degree of noise in experimental alkalinity measurements presents a challenge for accurate predictions [45].

Despite the overall high performance, the discrepancy in COD toward the end of the simulation suggests that the mass-based implementation may require further refinement in the particulate-to-soluble conversion ratios specifically for the later stage in the degradation of composite material. However, for the primary goals of operational stability tracking and gas production forecasting, the current mass-based ADM1 implementation provides a good reliable representation of the physical process.

#### 3.3.2 Laboratory Physical Twin R5

To evaluate the predictive reliability of the mass-based ADM1 beyond the initial calibration set, the model was validated using a co-reactor configuration. This involved applying the parameters calibrated from Physical Twin R6 to a second, independent system, Physical Twin R5, which featured distinct operational stressors and an extended timeframe.

While the calibration for R6 was conducted over 160 days, the validation for R5 was extended to 200 days to assess long-term model stability. A critical feature of the R5 configuration was a significant increase in the OLR between day 100 and day 150. This period served as a stress test for the model, forcing it to predict a rapid spike in metabolic activity and subsequent biogas production with results presented in Figure 7.

**Figure 7.**
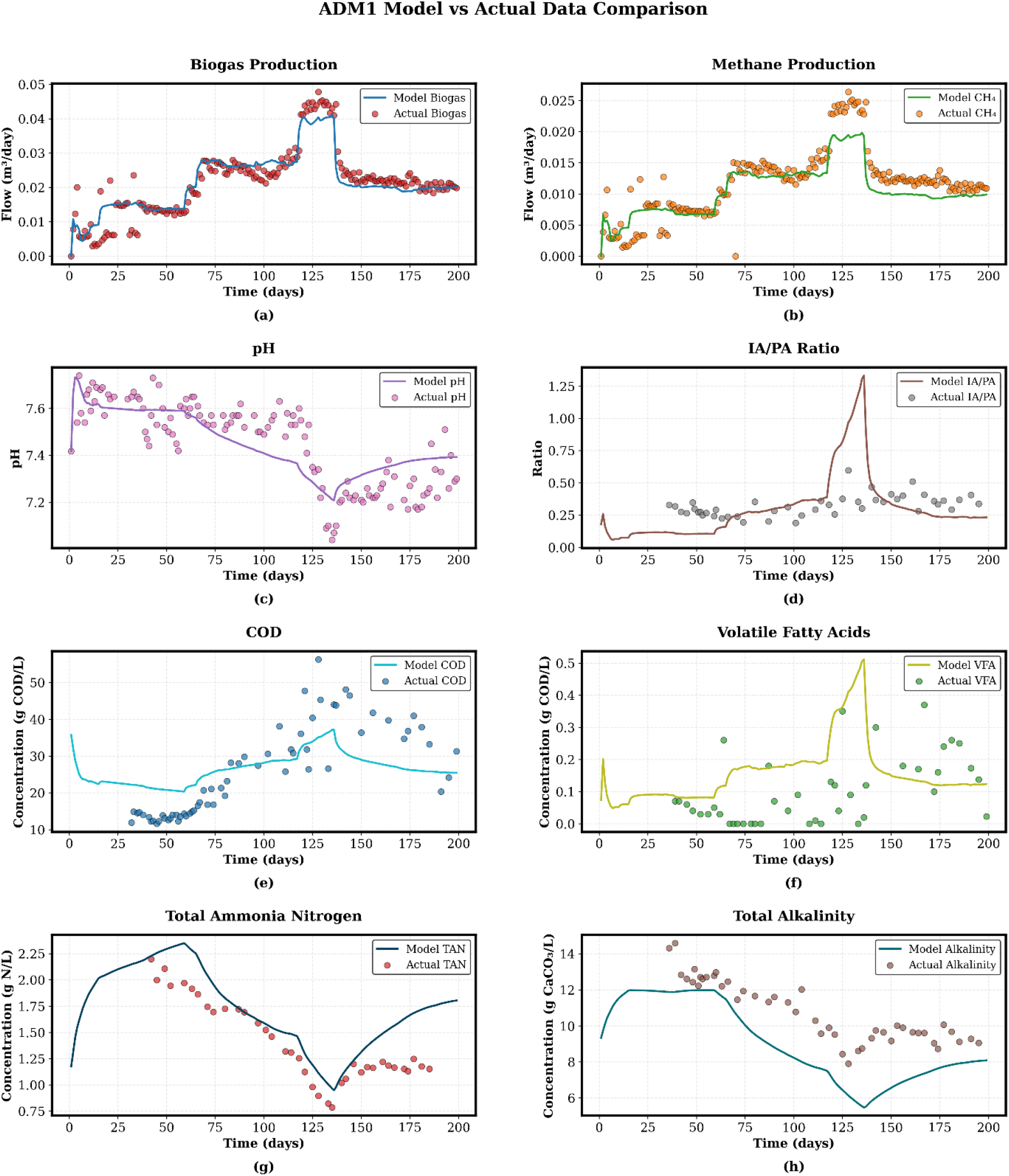
Simulation output and actual output comparison plots for key outputs of interest for physical twin reactor R5.

The model demonstrated superior predictive performance in the R5 co-reactor configuration compared to the R6 calibration, particularly regarding gas production and pH stability. The model successfully captured the biogas spike during the high-loading phase in R5, achieving a q_gas_ R^2^ of 0.85. This outperformed the R6 calibration, which recorded an R^2^ of 0.81 for the same variable. Similar improvements were noted in methane prediction q_ch4_, where the R^2^ increased from 0.71 in R6 to 0.74 in R5. The model’s ability to track the reactor’s acid-base balance improved in R5, with the R^2^ for pH rising to 0.62, compared to 0.38 in R6. This suggests that the mass-based framework is particularly effective at simulating stability over longer operational durations.

However, despite the overall high performance, certain variables showed consistent challenges across both configurations. Both R5 and R6 yielded negative R^2^ values for TAC and the IA/PA ratio. While the model captures the general downward trend in buffering capacity as OLR increases, the high volatility and sensitivity of experimental alkalinity measurements introduce significant noise that simulated values fail to match actual measured values. In R5, the COD reached concentrations exceeding 50 g COD/L during the peak loading phase. The model had a RMSE of 8.479 for COD in R5, slightly higher than the 7.33 observed in R6. This indicates that the mass-based conversion factors for complex particulates into soluble components may encounter limitations under extreme loading conditions. The successful validation on Physical Twin R5 confirms that the mass-based ADM1 is not overfitted to the calibration reactor. The improved performance metrics during the OLR spike in R5 demonstrate that the model accurately represents the underlying biochemical kinetics of the system, even when subjected to high-loading stress.

## 4.0 Conclusions

This study successfully developed, calibrated, and validated a mass-based ADM1 implementation tailored for the UK agricultural AD site. By applying a mass-based structure rather than the traditional COD-based framework, the model directly accommodates standard agricultural laboratory analysis, reducing the stoichiometric ambiguity typically associated with high-solids feedstocks like maize silage and cow manure.

The Global Sensitivity Analysis using the Sobol method proved to be a critical first step in the modelling workflow. By quantifying the total-order indices, the analysis identified k_dec_ as the globally dominant parameter, alongside key substrate-specific rates such as k_ch_ and k_m,ac_. This systematic identification allowed for a targeted calibration of just 10 to 13 high-impact parameters, preventing model over-parameterisation and ensuring that the final kinetic values remained physically meaningful.

The laboratory physical twin experimental design provided a high-fidelity data foundation that is often unattainable at industrial scales. The primary calibration on Physical Twin R6 yielded robust predictive accuracy for biogas production (R^2^ = 0.81) and methane yield (R^2^ = 0.71). By applying the R6-calibrated parameters to an independent replica, Physical Twin R5, the model demonstrated exceptional generalisability. It not only maintained high accuracy over an extended 200-day period but also successfully predicted the system’s dynamic response to a significant OLR stress test, achieving a biogas R^2^ of 0.85. The consistent performance across both physical twins confirms that a mass-based ADM1, when supported by a rigorous GSA and cross-reactor validation, serves as a reliable predictive tool at this scale. Building on these foundational results, future work will focus on validating this framework on a full-scale industrial reactor. This methodological framework offers the potential to open a scalable solution for the UK biogas industry, providing operators with a high-fidelity Digital Twin capable of forecasting process stability and optimising biogas production under the dynamic real-world conditions.

## CRediT authorship contribution statement

Rohit Murali: Writing – original draft, Investigation, Visualisation, Conceptualisation. Benaissa Dekhici: Writing – review & editing, Conceptualisation. Tao Chen: Writing – review & editing, Supervision, Project administration, Funding acquisition. Dongda Zhang: Writing – review & editing, Validation, Supervision, Project administration, Funding acquisition. Michael Short: Writing – review & editing, Supervision, Resources, Project administration, Investigation, Funding acquisition, Conceptualisation.

## Acknowledgements

This work was kindly supported and funded by the Surrey Institute for People-Centred AI (PAI) and by the Engineering and Physical Sciences Research Council EP/Y005600/1 (AI for Net Zero programme). The authors also thank the Supergen Bioenergy Impact Hub, EP/Y016300/1 for support.

