## Supplementary Material for "Modelling Anaerobic Co-Digestion with Agricultural Feedstock: Model Validation and Cross-Reactor Verification"

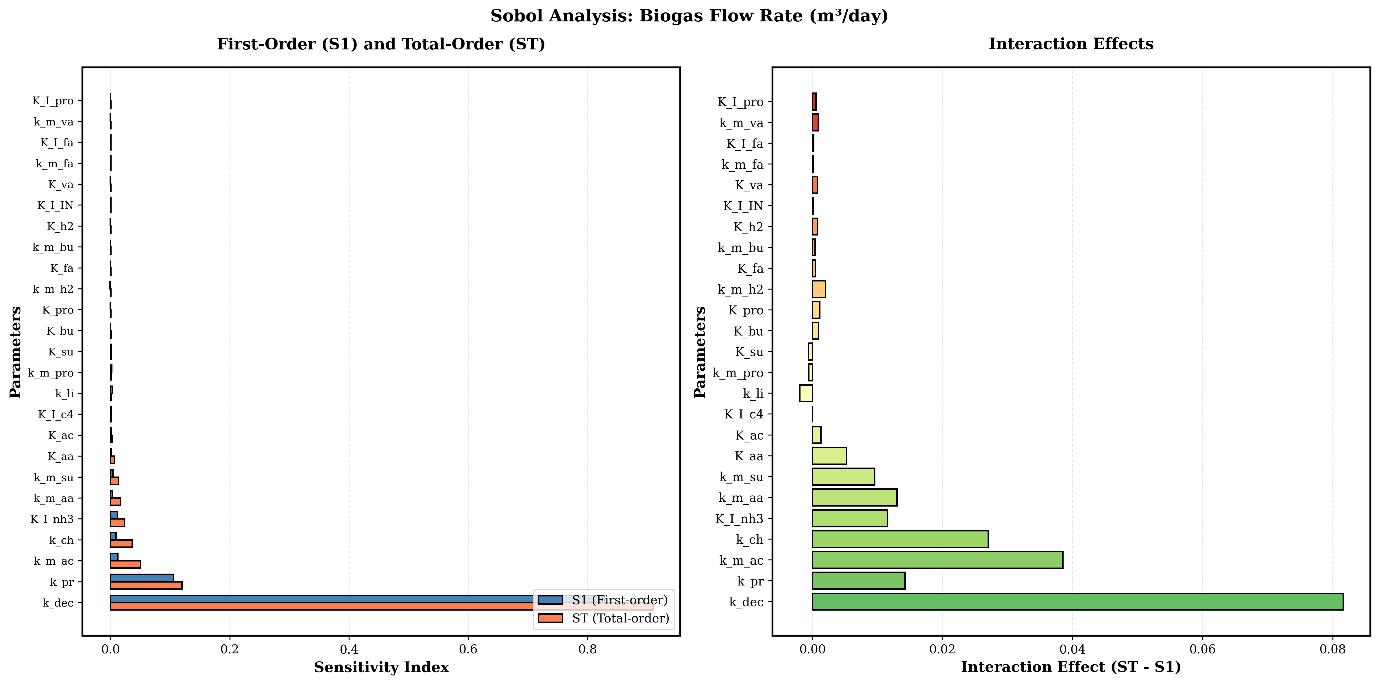

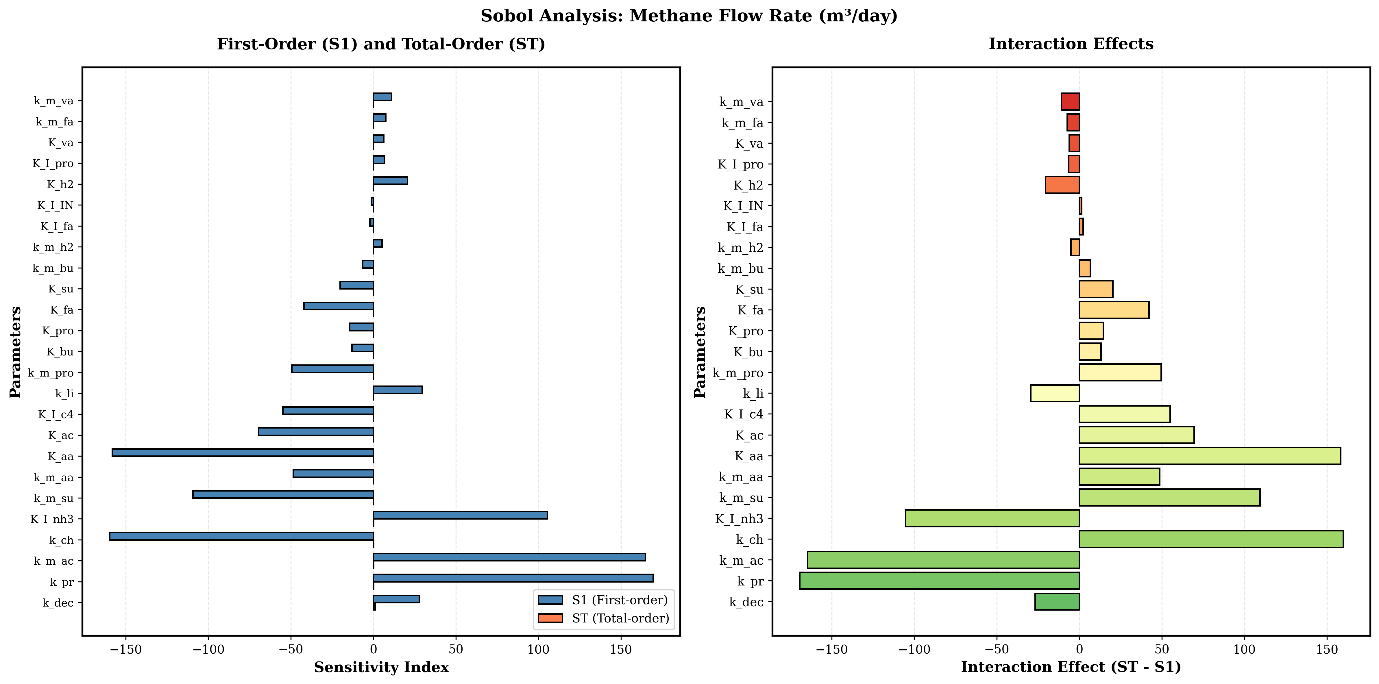
***Appendix A - Detailed Output from GSA***

Figure A1 - Sobol analysis of biogas flow rate for first order and total order sensitivity index on the left and interactions effects on the right.

Figure A2 - Sobol analysis of methane flow rate for first order and total order sensitivity index on the left and interactions effects on the right.

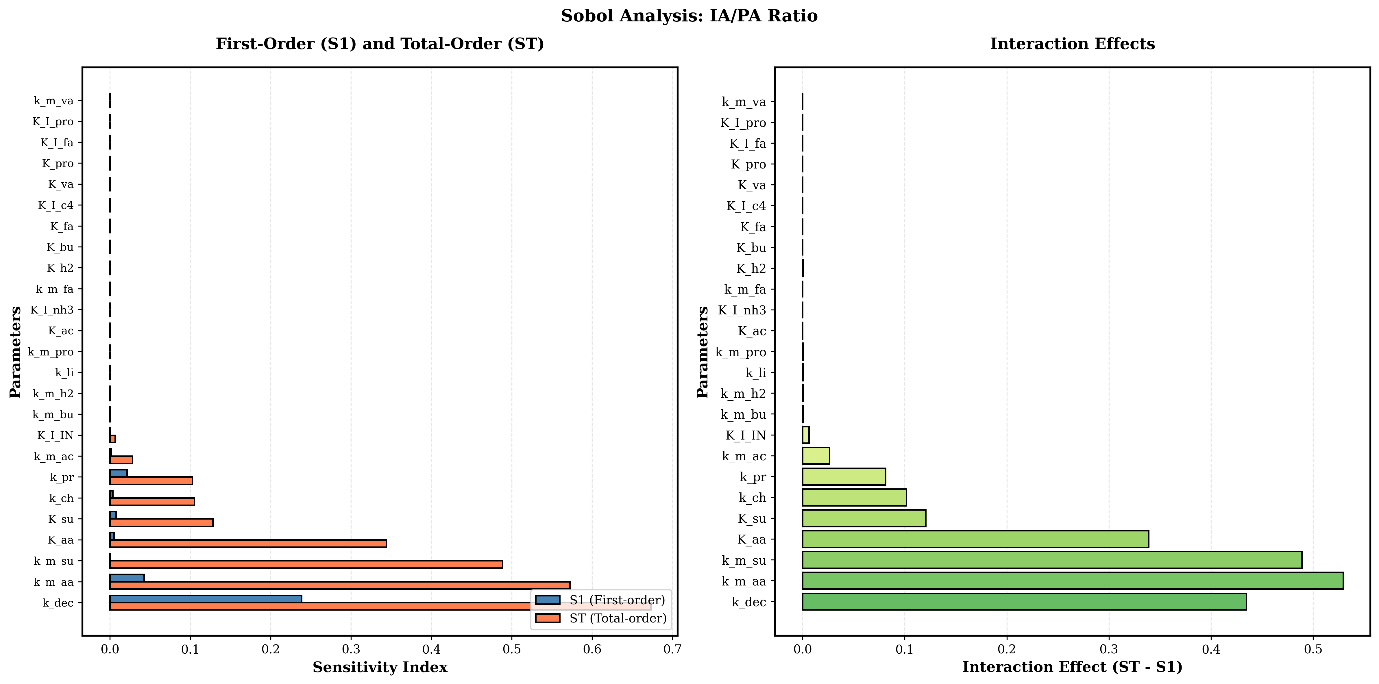

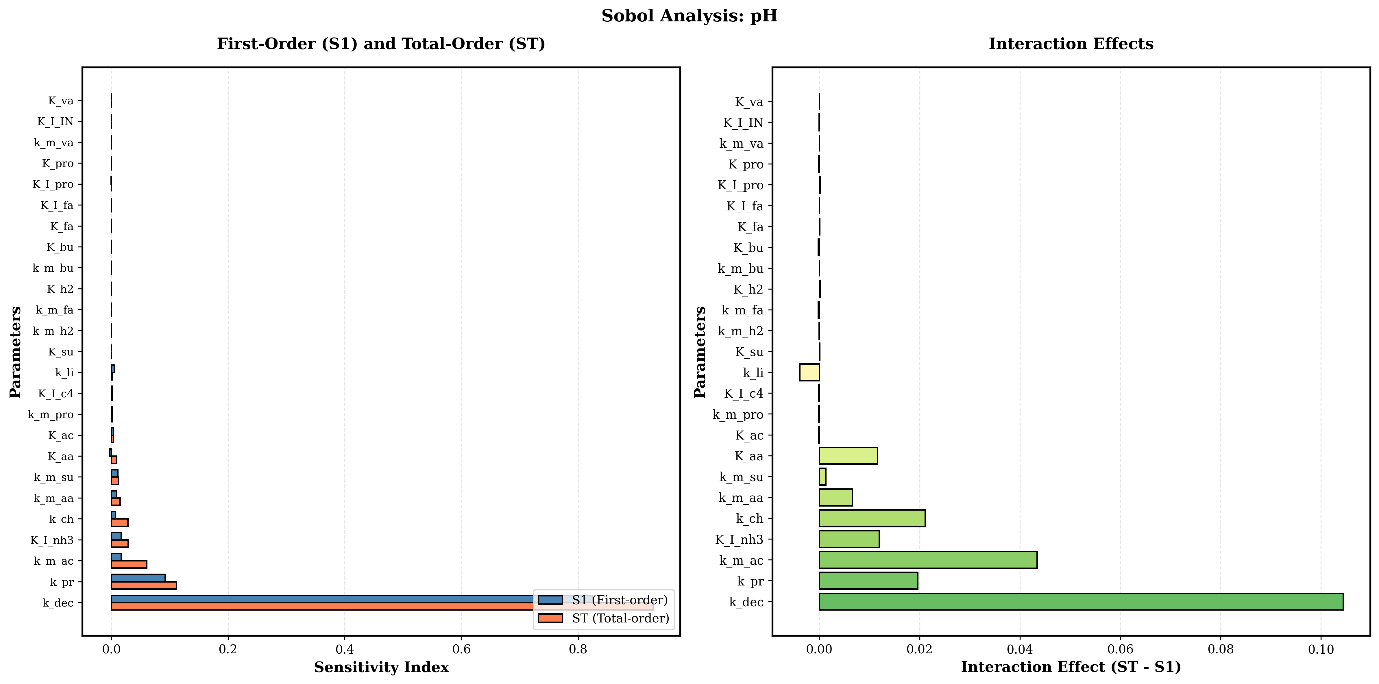

Figure A4 - Sobol analysis of IA/PA ratio for first order and total order sensitivity index on the left and interactions effects on the right.

Figure A3 - Sobol analysis of pH for first order and total order sensitivity index on the left and interactions effects on the right.

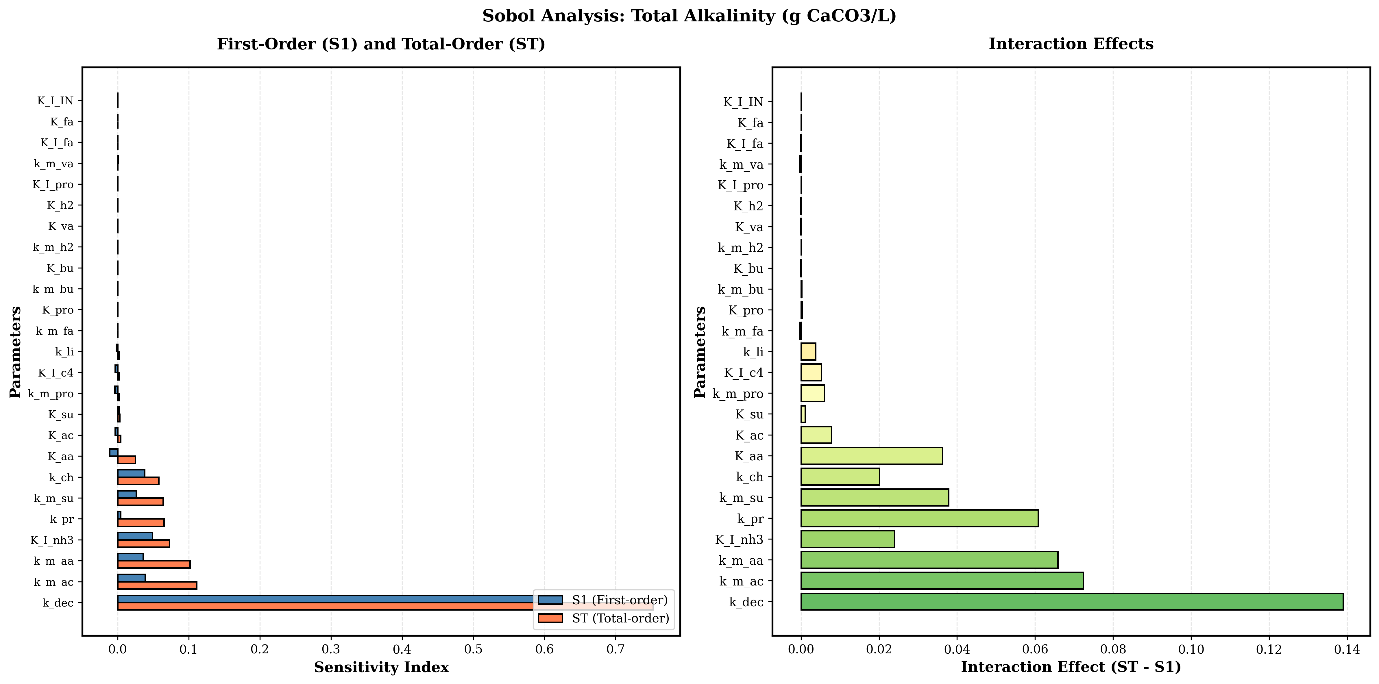

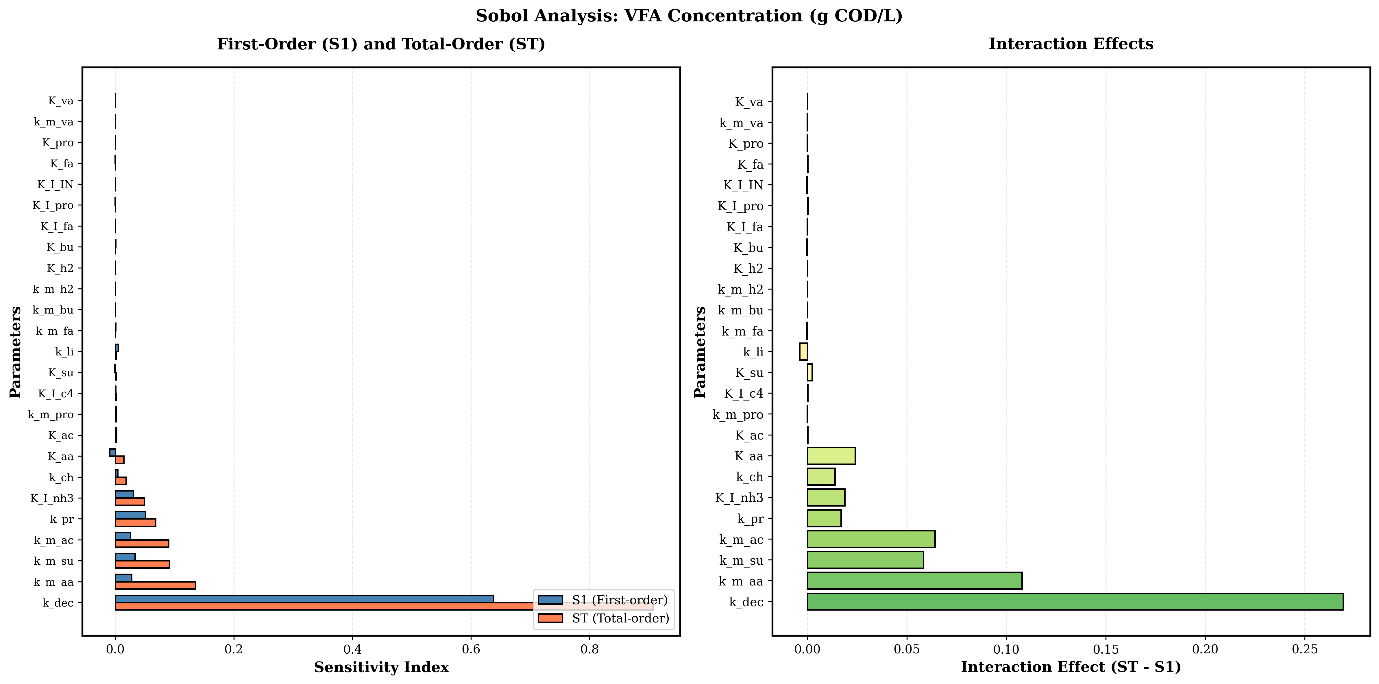

Figure A6 - Sobol analysis of total alkalinity for first order and total order sensitivity index on the left and interactions effects on the right.

Figure A5 - Sobol analysis of VFA concentration for first order and total order sensitivity index on the left and interactions effects on the right.

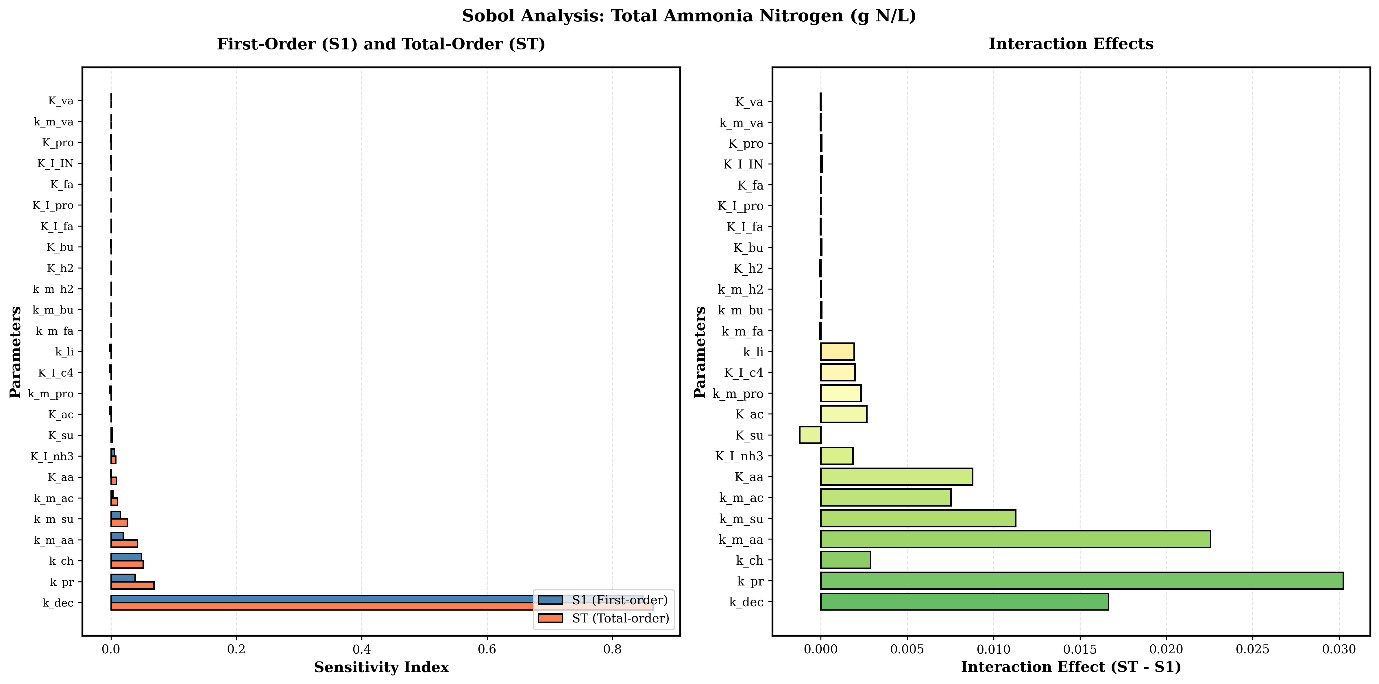

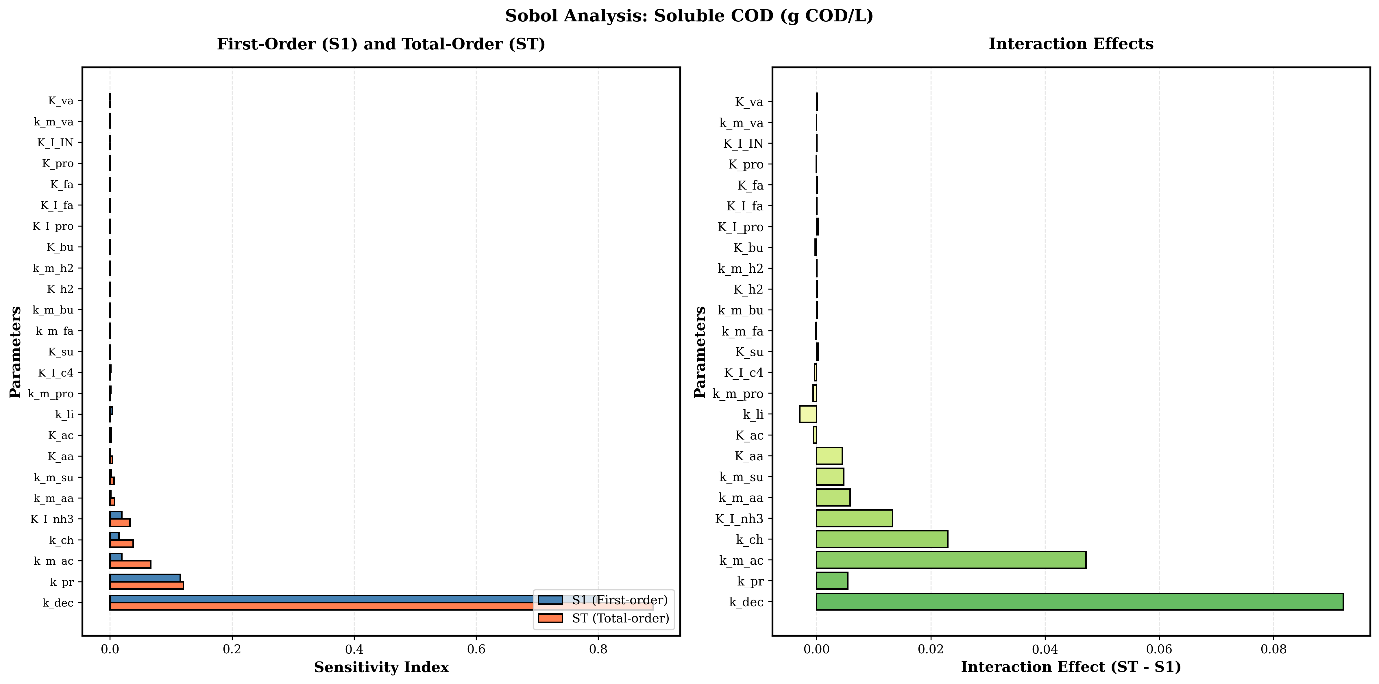

Figure A8 - Sobol analysis of total ammonia nitrogen for first order and total order sensitivity index on the left and interactions effects on the right.

Figure A7 - Sobol analysis of COD for first order and total order sensitivity index on the left and interactions effects on the right.
